# Integrated pretraining with evolutionary information to improve RNA secondary structure prediction

**DOI:** 10.1101/2022.01.27.478113

**Authors:** Zichao Yan, William Hamilton, Mathieu Blanchette

**Affiliations:** School of Computer Science, McGill University, Mila; School of Computer Science, Mila, McGill University

## Abstract

RNA secondary structure prediction is a fundamental task in computational and molecular biology. While machine learning approaches in this area have been shown to improve upon traditional RNA folding algorithms, performance remains limited for several reasons such as the small number of experimentally determined RNA structures and suboptimal use of evolutionary information. To address these challenges, we introduce a practical and effective pretraining strategy that enables learning from a larger set of RNA sequences with computationally predicted structures and in the meantime, tapping into the rich evolutionary information available in databases such as Rfam. Coupled with a flexible and scalable neural architecture that can navigate different learning scenarios while providing ease of integrating evolutionary information, our approach significantly improves upon state-of-the-art across a range of benchmarks, including both single sequence and alignment based structure prediction tasks, with particularly notable benefits on new, less well-studied RNA families. Our source code, data and packaged RNA secondary structure prediction software **RSSMFold** can be accessed at https://github.com/HarveyYan/RSSMFold.

## 1 Introduction

RNA secondary structure prediction is a fundamental task in computational and molecular biology. Given an RNA sequence *S*, the goal is to predict which pairs of nucleotides will form interactions (called base pairs) in the 3D structure *S* will fold into. Given the challenges of experimentally determining the structure of an RNA molecule, accurate computational structure prediction is essential for tasks such as RNA tertiary structure prediction Jonikas *et al*. (2009); Seetin and Mathews (2011), RNA-protein/ligand interaction Yan *et al*. (2020); Oliver *et al*. (2020) and therapeutic design of RNA aptamers Lieberman (2018); Bell *et al*. (2020).

Traditional approaches in this area have extensively relied on existing biological priors such as the decomposition of RNA secondary structures into nested substructures, as formulated in thermodynamic nearest neighbour models TINOCO *et al*. (1973); Pipas and McMahon (1975) and stochastic context free grammars (SCFG) Durbin *et al*. (1998); Do *et al*. (2006) based probabilistic models, which are amenable to exact dynamic programming algorithms McCaskill (1990); Zuker (2003); Lorenz *et al*. (2011). More recently, methods that leverage the power of representation learning have emerged, with learnt parameterized function that predicts RNA contact maps Singh *et al*. (2019); Chen *et al*. (2020).

Throughout these lines of works, the accuracy of the structure prediction pipeline has steadily improved, especially with the aid of more powerful deep learning techniques Singh *et al*. (2019), the combination of physics-based regularization with supervised learning Sato *et al*. (2021), and the use of evolutionary information extracted from multiple sequence alignment (MSA) Singh *et al*. (2021). However, several core challenges remain.

The first remarkable challenge is the limited amount of RNA secondary structure data. Compared to its protein counterpart, the task of predicting RNA secondary structures is systemically hindered by the lack of available training data. Among many standardized and publicly available RNA secondary structure benchmark datasets, the curated bprna dataset Danaee *et al*., 2018 is by far the largest one with 10,814 RNAs in the training set. Importantly, it carefully controls for sequence similarity within the training, validation and test sets. In comparison, another more recent dataset Chen *et al*. (2020), contains a significant amount of redundant sequences between validation and test set, and hence cannot be used to reliably measure model performance. Indeed, limiting sequence similarity to at most 80% reduces the dataset to approximately 3000 sequences. Second, evolutionary information is poorly exploited by deep learning models. While using multiple alignment of homologous sequences has long been known to increase predictive power Sankoff and Blanchette (1998), the best performing deep learning approach Singh *et al*. (2021) so far incorporating this type of information relies on external bioinformatics tools such as gremlin Kamisetty *et al*. (2013) to extract direct coupling signals. While such feature engineering strategy translates well on the benchmark, an end-to-end deep learning system operating directly on raw evolutionary data may improve performance and robustness.

In order to address the challenges outlined above, we introduce a flexible and scalable neural network architecture, referred as RNA Secondary Structure Model (**RSSM**), which we show achieves state-of-the-art performance across a range of benchmark tasks, including single-sequence structure prediction, evolutionary information aided structure prediction, and transfer learning from large-scale pretrained models using sequences and evolutionary information curated from the Rfam database Kalvari *et al*. (2020) along with computationally predicted structures.

Our contribution in this paper also includes outlining a general and practical strategy for improving RSSM generalization capability in new RNA families without any known structures, with particular benefits on pseudoknots and non-canonical basepairing interactions, and adapting RSSM to considerably longer RNA sequences with increased efficiency and accuracy through a sliding window based local structure prediction approach. We thoroughly analyze factors that can influence model performance, such as model ensembling and the impact of spurious homologs on alignment based RSSMs. We also investigate the robustness of alignment based RSSMs and improve its overall performance with the use of MSA transformer Rao *et al*. (2021).

## 2 Related work

RNA secondary structure prediction algorithms have seen a long line of development from the earlier physics-aided thermodynamic nearest neighbour models to more recent deep learning based approaches. Traditional RNA folding algorithms such as Mfold Zuker (2003), RNAstructures Reuter and Mathews (2010) and RNAfold Lorenz *et al*. (2011) are derived from the well-known Zucker’s algorithm Zuker and Stiegler (1981), which is based on dynamic programming that identifies the most stable RNA folding via minimizing an approximation to the molecule’s global free energy using experimentally determined thermodynamic parameters Lu *et al*. (2006); Turner and Mathews (2010). Nevertheless, due to the nature of nearest neighbour decomposition, these approaches generally ignore pseudoknots, a type of crucial basepairing interactions, although various heuristics have been proposed to circumvent this issue, most notably by exploiting the Boltzmann ensemble basepairing probabilities estimated from methods such as LinearPartition Zhang *et al*. (2020) and McCaskill’s algorithm McCaskill (1990). Representative approaches in this direction are IPknot Sato and Kato (2021) and ThreshKnot Zhang *et al*. (2019) that directly sample basepairs with high probabilities from a dense RNA contact map.

Learning-based methods represent another promising direction supported by the emergence of more RNA structure data. Early methods in this category are CONTRAfold Do *et al*. (2006); Andronescu *et al*. (2007), ContextFold Zakov *et al*. (2011) and Tornado Rivas *et al*. (2012), which combined the strength of traditional RNA folding algorithms with machine learning. Inductive bias such as substructure decomposition can be built into deep learning models, as demonstrated in MXFold2 Sato *et al*. (2021), although they come at the expense of higher computational complexity as well as the inability to identify pseudoknots. Other deep learning based methods such as SPOT-RNA Singh *et al*. (2019) and E2Efold Chen *et al*. (2020) directly predict RNA contact maps (matrix of basepairing probabilities) that naturally supports pseudoknots and any non-canonical interactions, although generalization to new RNA family that lies outside the scope of their training data is challenging Sato and Kato (2021).

Multiple sequence alignment (MSA) of evolutionarily related homologs can reveal co-variation patterns that are informative about interacting nucleotides and are used by approaches such as RNAalifold Bernhart *et al*. (2008) and CentroidAlifold Hamada *et al*. (2011). Direct coupling analysis Morcos *et al*. (2011); Kamisetty *et al*. (2013) (DCA) has also been used to identify potential interactions by capturing co-variation between MSA columns De Leonardis *et al*. (2015); Pucci *et al*. (2020). Such features are integrated into deep learning approaches such as SPOT-RNA2 Singh *et al*. (2021).

Recently, methods such as AlphaFold2 Jumper *et al*. (2021); Baek *et al*. (2021) for protein structure prediction have demonstrated that learning to extract and integrate evolutionary information can yield a dramatic increase in performance compared to their predecessors such as AlphaFold1 Senior *et al*. (2020). MSA transformer Rao *et al*. (2021) also demonstrates the superiority of learnt evolutionary features over DCA.

## 3 Task description

Figure 1 illustrates four types of benchmark tasks considered in our study, which are single sequence based and multiple sequence alignment based RNA secondary structure prediction, either with pretraining or trained from scratch. We detail the setup for our benchmark tasks below.

**Figure 1:**
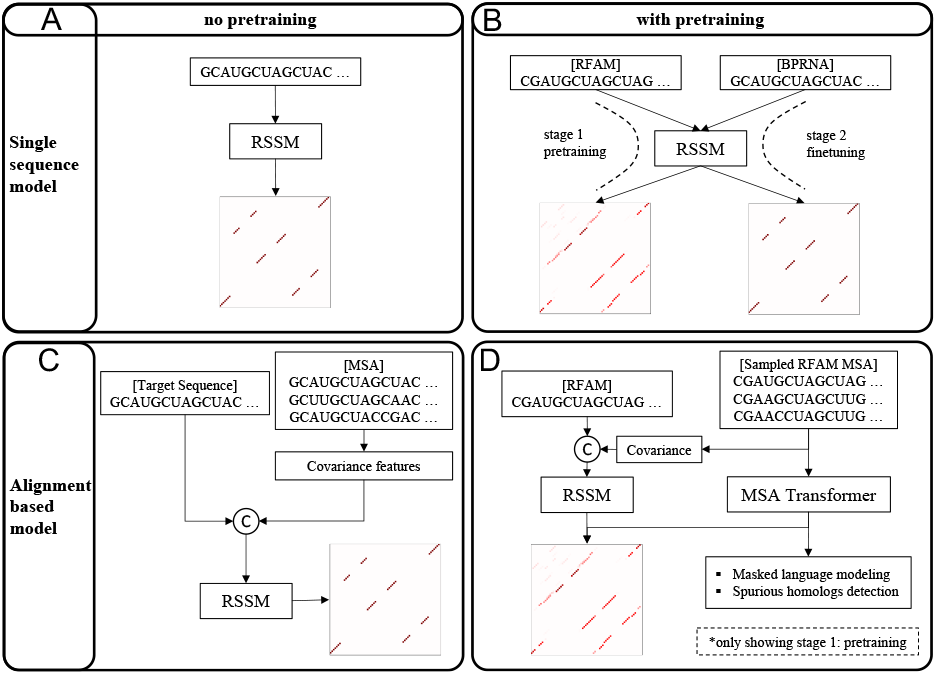
An illustration of the four types of tasks used in our study, with or without pre-training, and with or without using homologous sequences. (A) Single sequence based RSSM learns to predict RNA secondary structure on a per RNA basis in the bprna dataset. (B) RSSM is pretrained using RNAs selected from Rfam database with computationally predicted structures, then finetuned in the bprna dataset. (C) Evolutionary information in the form of covariance matrix can be incorporated into RSSM via concatenation with the initial RNA contact map representation. (D) The Rfam database also provided manually curated alignments which we seek to integrate into the pretraining process, either in its extracted covariance feature form or through a dedicated MSA transformer.

### 3.1 Training single sequence based models

While adding evolutionary information usually improves RNA structure prediction, acquiring such information from gigantic nucleotide database and feeding them to the model in a desirable format often generate large computational pressure that undermines the efficiency of the whole predictive pipeline Zhang *et al*. (2021); Singh *et al*. (2021). The availability of evolutionary information is also heavily influenced by factors such as the extent to which the RNA in question is conserved and whether its homologs can be retrieved from the database. Therefore, as the first task considered in our study (Figure 1 A), we train single sequence based RSSMs using the same bprna dataset originally curated by Singh *et al*., 2019, which contains 10,814 sequences in the training set (TR0), 1300 sequences in the validation set (VL0) and 1305 sequences in the test set (TS0). RNA sequences in bprna dataset only go up to 500 nts. We also evaluate our RSSMs on a smaller PDB (67 sequences; TS1) and NMR (39 sequences; TS2) dataset ^2^ featuring shorter RNA sequences (33-189 nts), which are also curated by the same authors Singh *et al*. (2019) and had its sequence similarity controlled at 80% against bprna dataset.

In addition, we use an external dataset from Sato and Kato, 2021 to evaluate the generalization capability of our RSSMs in new RNA families and longer RNAs up to 4381 nts, with sequence similarity controlled at 80% against bprna dataset. We further remove sequences above 80% similarity from the external dataset against our own pretraining data. More details about this external dataset is in Suppl. section B.

### 3.2 Training alignment based models

In this task (Figure 1 C), we consider providing evolutionary information to RSSM. For this purpose, we use the RNAcmap tool Zhang *et al*. (2021) to obtain MSAs from the NCBI nucleotide database (shown in Figure 2 right), for each RNA sequence of interest. Through the initial seed alignment returned by blastn Madden (2013) and annotated consensus secondary structure (CSS), RNAcmap builds a covariance model Durbin *et al*. (1998) for the target sequence and employs Infernal Nawrocki and Eddy (2013) to scan the nucleotide database again to obtain the final MSA. In particular, an accurate CSS potentially reveals evolutionarily co-conserved RNA basepairs, which helps the subsequent Infernal stage to discover more related RNA homologs. For the annotation of CSS inside the RNAcmap pipeline, we use either our own single sequence based RSSM or RNAfold Lorenz *et al*. (2011), considering the recent findings in Sato and Kato, 2021 that thermodynamic models such as RNAfold tend to generalize better to new Rfam families compared to deep learning based models such as SPOT-RNA.

**Figure 2:**
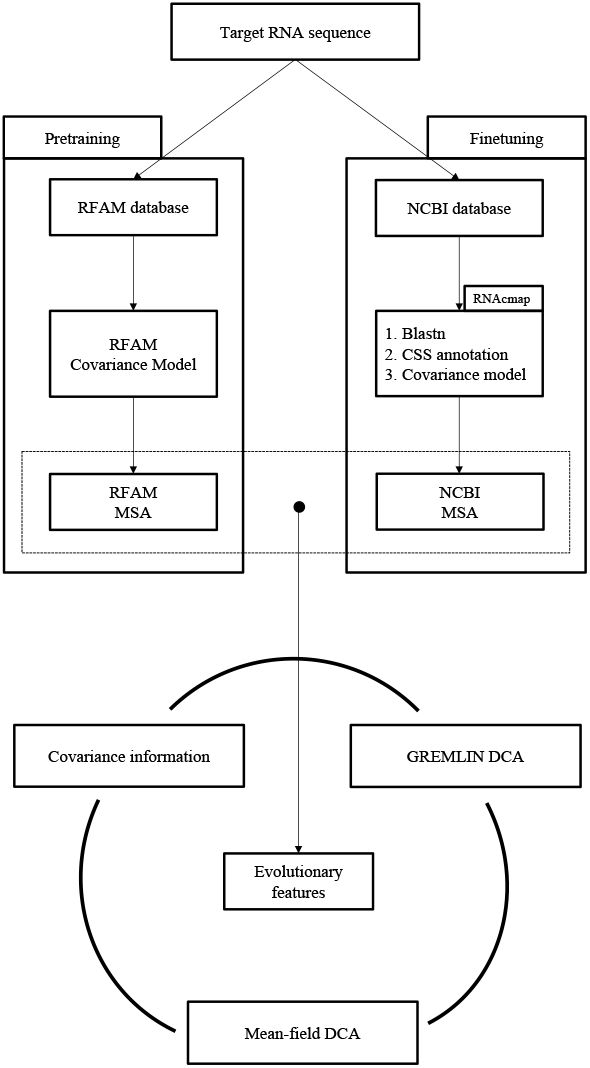
Our study considers two approaches for obtaining multiple sequence alignment. Approach on the left uses Rfam deposited covariance models to classify and align target RNAs to existing RNA families in the Rfam database. Despite its simplicity, we emphasize that this method cannot determine structures for RNAs from unknown RNA families. Nevertheless, we have used this method to obtain Rfam MSAs for evolutionary information integrated pretraining. The other approach shown on the right, having the RNAcmap tool at its core, is able to construct MSAs for RNAs from any unknown RNA families.

We consider two approaches for exploiting and integrating evolutionary information from MSAs into our alignment based RSSMs, briefly summarized below and to be substantially detailed in the Methods section:

- Extracting covariance features from MSAs by gathering MSA single and pairwise column statistics. It is worth pointing out that covariance matrix of MSA has formed the basis of various direct coupling analysis pipelines such as mean-field DCA Morcos *et al*. (2011).
- End-to-end learning of MSAs through a hybrid RSSM which includes a MSA transformer Rao *et al*. (2021) component.

Similar to the single sequence based RSSMs, we also train our alignment based models on the bprna dataset and evaluate them on the external dataset from Sato and Kato, 2021.

### 3.3 Transfer learning from Rfam pretrained models

To overcome the limitation of training data, we consider supervised pretraining for our single sequence as well as alignment based RSSMs using only RNA sequences in the Rfam database Kalvari *et al*. (2020) (v14.5), as shown in Figure 1 B and D. Here, we leverage non-coding RNA sequences from Rfam database as they are mostly associated with biologically meaningful regulatory functions that are useful to other studies. The Rfam database also provides manually curated alignments that can be used later for our evolution-based pretraining.

We create our pretraining RNA sequence library by combining sequences shorter than 1000 nts across all Rfam families, followed by removing redundant sequence. In particular, cd-hit-est Fu *et al*. (2012) is used to control the sequence similarity within the Rfam sequence library at 80%, and cd-hit-est-2d is used to remove sequences which share at least 80% similarity with sequences in the bprna test set. The 80% sequence identity is the lowest allowed cutoff by cd-hit-est and it has been used similarly in SPOT-RNA Singh *et al*. (2019) and MXFold2 Sato *et al*. (2021).

We end up with a set of 184,226 unique RNA sequences, for which we also obtain computationally predicted structures in the form of ensemble basepairing probabilities from LinearPartition Zhang *et al*. (2020), a state-of-the-art thermodynamic RNA folding model Zhang *et al*. (2019); Sato and Kato (2021). By leveraging LinearPartition’s rich predictions, our supervised pretraining strategy is applicable to any RNA sequences without requiring the knowledge of their actual structures.

For single sequence based pretraining, we perform basepairing probability prediction and non-coding RNA classification. For evolutionary information integrated pretraining, sequences belonging to each RNA family are aligned (Figure 2 left), and produces a subsampled MSA which is purposefully contaminated with spurious homologs each time an RNA sequence in sampled from the library. Afterwards, we either manually extract covariance features to pretrain our RSSM or employ a MSA transformer Rao *et al*. (2021) to learn to extract evolutionary information from MSAs. We elaborate more in section “Pretraining strategies”.

## 4 Methods

The overall architecture of RSSM is shown in Figure 3 A. RSSM reflects our design principle to balance the focus on short and long range RNA secondary structure basepairs, which motivates the use of a hybrid architecture consisting of dedicated local and global modules. Indeed, we have observed that transformer-only Vaswani *et al*. (2017) architectures can be subject to severe overfitting (Suppl. Figure S9).

**Figure 3:**
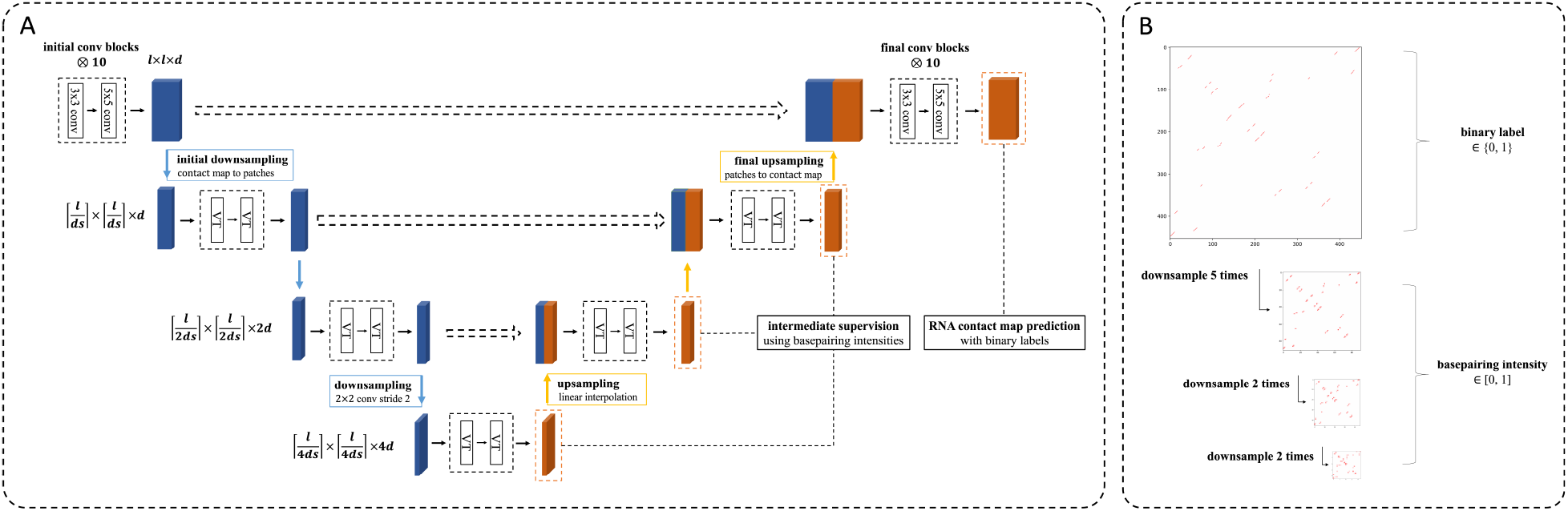
(A) Our model is designed with the principle to balance its focus on local and long-range RNA basepairing interactions. For this purpose, RSSM adopts a u-net architecture, with a convolutional backbone at the top dedicated to local RNA basepairs and vision transformers operating on spatially downsampled RNA feature maps for the extraction of longer range interactions. (B) RSSM can be trained with intermediate supervision in the form of basepairing intensity scores. Refer to Suppl. section A for more details.

The input to our RSSMs are outer-concatenated RNA sequence embedding yielding an ℝ^*l*×*l*×*d*^ representation, with *l* being the length of RNA and *d* denoting the base dimension of RSSM. In alignment based RSSMs, outer-concatenated one-hot encoding of the RNA sequence are used instead, and upon concatenation with evolutionary information, undergoes a 3 × 3 convolution layer to obtain the initial RNA contact map representation.

### 4.1 RSSM convolutional backbone

Based on the above consideration, our model resorts to a u-net architecture Ronneberger *et al*. (2015), with convolutional modules LeCun *et al*. (1989) at the top operating on the full RNA contact map representation. We arrange our convolutional modules into blocks, each consisting of a 3 × 3 convolution followed by a 5 × 5 convolution. Each convolutional layer is preceded by layer normalization Ba *et al*. (2016), activation function and dropout Srivastava *et al*. (2014) along with residual connection He *et al*. (2016).

Our RSSM contains 20 such blocks and they are considered the backbone of our model, as feature maps generated by the first half of these convolutions are used as input to the later self-attention modules, while the latter half produce the final RNA contact map representation.

### 4.2 Self-attention on spatially downsampled RNA contact map representation

For a given RNA contact map representation *M* ∈ ℝ^*l*×*l*×*d*^, an ordinary self-attention proceeds by first flattening the map along its spatial dimensions into 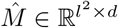. Then, self-attention proceeds as follows:

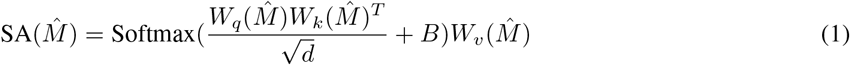

where *W*_*q*_, *W*_*k*_ and *W*_*v*_ respectively map query, key and value vectors from 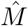, and *B* represents a bias matrix which can be used to encode positional information in *M*.

Despite its straightforwardness, the computational complexity in Eq. 1 is *𝒪* (*l*^4^), which poses considerable optimization challenge for RNAs of a few hundred nucleotides or more. In order to facilitate such computation, we implement two strategies: (i) downsampling the original spatial dimensionality of *M*, and (ii) limiting the scope of self-attention.

Below the convolutional backbone, our model looks at spatially downsampled RNA feature maps, with dimension being 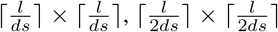 and 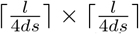. This follows the approach of a vision transformers Dosovitskiy *et al*. (2021) (denoted by VT). These downsampled maps are obtained using convolution layers with strides of *ds*, 2 and 2, which merge spatially contiguous regions while doubling the number of feature channels. Self-attention is only applied to these downsampled feature maps, which incurs less computational overhead while simultaneously facilitating the discovery of longer range interactions.

These downsampled feature maps are connected to the original RNA contact map representation through the horizontal shortcut and expansive path in the u-net architecture, which allows spatially condensed information from lower levels to be gradually merged into the RSSM backbone. In each upsampling step, we concatenate linearly interpolated lower level features with a corresponding map from the shortcut, which then undergo a 3 × 3 convolution layer followed by another two vision transformer modules. Along the contractive path, we insert paddings whenever necessary. Similarly, linear interpolated feature maps are cropped to align the corresponding downsampled map from the contractive path.

To limit the scope of self-attention, we followed the local window approach in swin-transformer Liu *et al*. (2021), which confines the computation of self-attention within non-overlapping local windows. Assuming a window size of *w*, RNA contact map representation *M* is partitioned into non-overlapping patches of size *w* × *w* which leads to 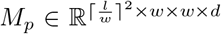, and self-attention simply computes within each patch. Using this localized mode of selfattention, the computational complexity can be brought down to *𝒪* (*w*^2^*l*^2^) which is more practical with longer RNAs. In addition, we apply shifted local windows for each consecutive transformer module as well as relative positional embeddings in Eq. 1.

### 4.3 Integrating evolutionary information into RSSM

Given a MSA, we first consider manually extracting covariance features, which involves gathering single MSA column and pairwise columns statistics under a sequence re-weighting scheme meant to correct for phylogeny sampling bias. We resort to the same calculation as Morcos *et al*., 2011 (refer to Eq.1, Eq.2 and Eq.10). Here, an analogy can be drawn between covariance features for MSAs and one-hot encodings commonly used for single sequences. We denote the extracted covariance matrix by 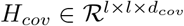.

Alternatively, we feed MSA directly to MSA transformer, producing learnt MSA embedding *h* ∈ ℝ^*m*×*l*×*d*^, and all row-wise attention weights *a* ∈ ℝ^*l,l,d′*^. Here, *m* and *l* denote depth and length of the MSA, and the final extracted evolutionary features are in the form: [*H*_*tt*_ || *a*], where *H*_*tt*_ is the outer concatenation of *h*_*t*_, representation of the target RNA sequence with its index in the MSA denoted by *t*, and [ …. || … ] denotes concatenation along the feature dimension.

Given either *H*_*cov*_ or [*H*_*tt*_ || *a*], RSSM concatenates evolutionary information with outer-concatenated RNA sequence one-hot encoding along the feature dimension, then uses a 3×3 convolution to obtain the initial RNA contact map representation.

### 4.4 Pretraining strategies

The core objective of pretraining is to overcome the limited size of our labeled training set, which otherwise poses considerable challenge to the generalization capability of our RSSMs in new RNA families and longer RNAs.

#### For single sequence pretraining

we leverage predicted basepairing probabilities from LinearPartition Zhang *et al*. (2020) as learning targets in place of the ordinary ground truth binary labels (Eq. 3). It is worth emphasizing again that one of the major advantages of our supervised pretraining strategy is its applicability to any RNA sequences without requiring knowledge of their actual structures.

In addition, we consider non-coding RNA classification Childs *et al*. (2009); Panwar *et al*. (2014), motivated by the fact that RNAs in a conserved family share similar structures, as a consequence of evolution. Therefore, non-coding RNA classification in our multitask pretraining paradigm may provide additional supervision signals to help RSSMs identify distinct domains of RNA structures.

We classify non-coding RNAs by their superfamilies according to RNArchitecture Boccaletto *et al*. (2017). A total of 91 superfamilies are selected, each represented by at least 100 RNA sequences in our curated pretraining dataset, with some of the larger superfamilies being transfer RNA (tRNA), C/D-box RNA, large subunit ribosomal RNA (LSU rRNA) and Cobalamin riboswitch, each containing more than 7000 sequences. Sequences from less well-represented RNA superfamilies are excluded from this type of supervision.

To demonstrate the effect of non-coding RNA classification, we present in Figure S1 projections of RNA embeddings to two dimensional space using t-SNE van der Maaten and Hinton (2008), along with RNA secondary structures predicted for a few representative RNA families.

#### In alignment based pretraining

for each RNA sequence we sample a subset of its homologs from our curated pretraining sequence library, and align them into a MSA with the corresponding Rfam covariance model (Figure 2 A). In particular, we add spurious homologs to these subsampled Rfam MSAs, in the hope of accommodating the potential quality discrepancies between Rfam’s expert annotated MSAs and ad-hoc MSAs obtained by the RNAcmap pipeline in other dataset. To this end, 10% of sequences in Rfam subsampled MSAs are manually contaminated with spurious homologs, with half of these sequences randomly replacing 10% of their nucleotides with { *A, C, G, U*, − }, and the second half having 20% of their *k*-mers replaced by contents in other RNA families (*k* being a uniformly sampled integer from 5 to 15).

We either directly incorporate evolutionary information from MSAs into RSSM via extraction of covariance matrices from the entire subsampled MSAs, or in an end-to-end manner with MSA transformer Rao *et al*. (2021), as we have shown in the “Integrating evolutionary information into RSSM” section.

In addition to basepairing probability prediction, our MSA transformer is pretrained with masked language modeling (MLM) objective Devlin *et al*. (2019), with 15% of tokens in the non-noisy part of the MSAs randomly masked, as well as spurious homolog detection. Given a learnt MSA *h* ∈ ℝ^*m*×*l*×*d*^ and all row-wise attention weights *a* ∈ ℝ^*l,l,d′*^, the overall pretraining objective can be expressed as follows:

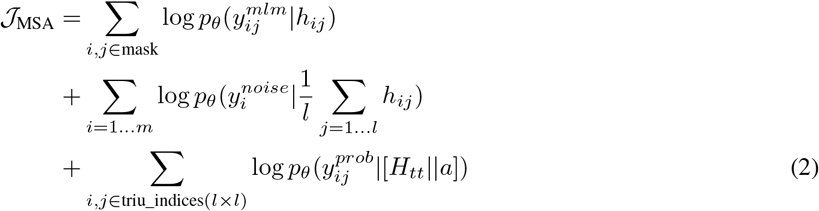

Concerning our computational budget, we keep the maximum depth of randomly subsampled MSA at *m* = 2^15^*/l* during the course of pretraining. We make sure that the target sequence is always included in subsampled MSAs.

## 5 Results

Information on hyperparameters and training details is in Suppl. section A. Here, we only present our core results.

### 5.1 Performance of single-sequence based models

We present the performance for our single-sequence based RSSM in Table 2, along with a variety of baselines on the bprna test set. All our models are trained 300 epochs using the bprna training set and we are able to pinpoint several factors that benefit RSSM performance. For each base dimension and transfer learning setting, we train five RSSM replicates with random weight initialization and register the mean and standard deviation of their performance under single models.

First of all, doubling the base dimension of RSSM from 64 to 128 yields a consistent 4% F1 improvement across most training setting. Without doubt, these gains come at a cost in terms of higher computational demand on GPU hardware and training time.

Pretraining using Rfam significantly improves performance across various settings. It helps maintaining a reasonable computational overhead with *d* = 64, while achieving nearly comparable performance to RSSM with higher hidden dimension *d* = 128. Furthermore, utilizing our curated pretraining data allows RSSM to achieve better generalization capability in new RNA families absent from the bprna finetuning dataset, which we will elucidate later in section “Generalization to new RNA families” and Suppl. section B.

Model ensembling by simply averaging the predicted contact map basepairing probabilities across the five replicates also substantially improves performance. Although all replicates share the same neural architecture, random initialization and shuffled training help each model learn different hypotheses, and ensemble averaging serves as a mean of trimming outlier predictions while retaining confident basepairs that all replicates agree on. We look into this phenomenon in Suppl. section F. Indeed, this simple ensembling strategy has proven to be essential later in our attempt to boost RSSM generalization capability in new RNA families (section “Performance of alignment based models”) and in our end-to-end alignment based RSSM (section “Performance of alignment based models”).

Our RSSMs have shown consistent performance gains compared to thermodynamic baselines such as RNAfold, CONTRAfold and LinearPartition based methods, in addition to a deep learning based model that is SPOT-RNA. We point out that SPOT-RNA is also a five-model ensemble and is built on convolutional neural network architectures, to which our RSSM convolutional backbone shares similar design.

Furthermore, we seek to measure model performance on different classes of basepairs, based on the distance between the two interacting nucleotides along the RNA sequence. To this end, we classify all RNA basepairs in the bprna test set into three groups that represent short, medium and long range interactions, and our results are shown in Table 1.

**Table 1:** Performance of our single sequence based RSSMs and baselines on three groups of bprna test set basepairs, which are separated by the distance between two basepairing nucleotides along the RNA sequence (e.g. below 200 nts, between 200 and 400 nts, and above 400 nts). The number of such basepairs is indicated inside parentheses.

| Mode | Model | < 200 nts (support: 38856 bps) |  |  | 200~400 nts (support: 1848 bps) |  |  | ≥ 400 nts (support: 81 bps) |  |  |
| --- | --- | --- | --- | --- | --- | --- | --- | --- | --- | --- |
|  |  | F1 | Precision | Recall | F1 | Precision | Recall | F1 | Precision | Recall |
| Single models | RSSM<sub>d64</sub> | 0.649±0.002 | 0.681±0.008 | 0.620±0.007 | 0.707±0.008 | 0.763±0.021 | 0.660±0.004 | 0.714±0.024 | 0.827±0.039 | 0.630±0.027 |
|  | RSSM<sub>d64,T</sub> | 0.671±0.003 | 0.699±0.006 | 0.645±0.005 | 0.739±0.005 | 0.775±0.006 | 0.706±0.006 | 0.686±0.017 | 0.825±0.039 | 0.588±0.013 |
|  | RSSM<sub>d128</sub> | 0.672±0.002 | 0.704±0.010 | 0.643±0.010 | 0.719±0.007 | 0.774±0.012 | 0.672±0.019 | 0.743±0.030 | 0.847±0.056 | 0.664±0.029 |
|  | RSSM<sub>d128,T</sub> | 0.701±0.001 | 0.737±0.014 | 0.668±0.011 | 0.744±0.005 | 0.785±0.020 | 0.708±0.012 | 0.710±0.016 | 0.791±0.045 | 0.644±0.009 |
| Ensemble models | RSSM<sub>d64</sub> | 0.671 | 0.722 | 0.626 | 0.725 | 0.808 | 0.657 | 0.739 | 0.895 | 0.630 |
|  | RSSM<sub>d64,T</sub> | 0.686 | 0.725 | 0.651 | 0.745 | 0.788 | 0.706 | 0.706 | 0.873 | 0.593 |
|  | RSSM<sub>d128</sub> | 0.701 | 0.757 | 0.652 | 0.754 | 0.838 | 0.685 | 0.786 | 0.932 | 0.679 |
|  | RSSM<sub>d128,T</sub> | 0.725 | 0.776 | 0.680 | 0.757 | 0.806 | 0.713 | 0.738 | 0.867 | 0.64 |
| Baselines | SPOT-RNA | 0.588 | 0.583 | 0.594 | 0.509 | 0.656 | 0.416 | 0.653 | 0.605 | 0.710 |
|  | RNAfold | 0.485 | 0.423 | 0.569 | 0.404 | 0.390 | 0.418 | 0.480 | 0.403 | 0.593 |
|  | CONTRAfold | 0.521 | 0.465 | 0.593 | 0.453 | 0.477 | 0.431 | 0.571 | 0.575 | 0.568 |
|  | IPknot |  |  |  |  |  |  |  |  |  |
|  | LinearPartition-C | 0.526 | 0.491 | 0.566 | 0.265 | 0.429 | 0.192 | 0.296 | 0.501 | 0.210 |
|  | LinearPartition-V | 0.494 | 0.428 | 0.583 | 0.408 | 0.399 | 0.417 | 0.490 | 0.407 | 0.617 |
|  | ThreshKnot |  |  |  |  |  |  |  |  |  |
|  | LinearPartition-C | 0.540 | 0.540 | 0.540 | 0.249 | 0.174 | 0.435 | 0.301 | 0.531 | 0.210 |
|  | LinearPartition-V | 0.495 | 0.432 | 0.581 | 0.405 | 0.399 | 0.412 | 0.498 | 0.417 | 0.617 |

Once again, we observe the benefits of larger base dimension, model ensembling and pretraining across all basepair interaction ranges, except pretrained models which do not contribute to the accuracy of longer range basepair prediction, possibly due to the lack of useful long range basepairing signals in our computationally obtained structures. Nevertheless, our RSSMs have maintained balanced performance among short, medium and long range basepairs, and outperformed all other baselines. In particular, comparing RSSM_d64_ to the pure convolutional baseline i.e. SPOT-RNA suggests the benefit of our attention based modules.

### 5.2 Generalization to new RNA families

All our RSSM variants in Table 2 significantly outperform the baselines as they have leveraged the power of representation learning. Traditional RNA folding algorithms such as RNAfold, CONTRAfold and LinearPartition based methods have built-in inductive bias such as the decomposition formula for large RNA secondary structures. In comparison, deep learning models such as SPOT-RNA and RSSMs may fail to generalize to new RNA families outside the scope of training data.

**Table 2:** Performance of our single sequence based RSSMs and a variety of baselines, on the bprna dataset. Subscripts indicate the base dimension (d64 or d128) of RSSM and use of transfer learning (T).

| Mode | Model | Test set |  |  |
| --- | --- | --- | --- | --- |
|  |  | F1 | Precision | Recall |
| Single models | $\text{RSSM}_{d64}$ | $0.656 \pm 0.002$ | $0.662 \pm 0.006$ | $0.674 \pm 0.006$ |
| | $\text{RSSM}_{d64,T}$ | $0.671 \pm 0.003$ | $0.689 \pm 0.005$ | $0.676 \pm 0.005$ |
| | $\text{RSSM}_{d128}$ | $0.679 \pm 0.003$ | $0.700 \pm 0.005$ | $0.682 \pm 0.008$ |
| | $\text{RSSM}_{d128,T}$ | $0.702 \pm 0.001$ | $0.728 \pm 0.011$ | $0.698 \pm 0.008$ |
| Ensemble models | $\text{RSSM}_{d64}$ | 0.672 | 0.668 | 0.704 |
| | $\text{RSSM}_{d64,T}$ | 0.683 | 0.715 | 0.683 |
| | $\text{RSSM}_{d128}$ | 0.700 | 0.737 | 0.690 |
| | $\text{RSSM}_{d128,T}$ | 0.720 | 0.758 | 0.708 |
| Baselines | SPOT-RNA | 0.626 | 0.709 | 0.560 |
|  | RNAfold | 0.491 | 0.449 | 0.577 |
|  | CONTRAfold | 0.522 | 0.484 | 0.598 |
|  | IPknot |  |  |  |
|  | LinearPartition-C | 0.525 | 0.514 | 0.569 |
|  | LinearPartition-V | 0.500 | 0.455 | 0.591 |
|  | ThreshKnot |  |  |  |
|  | LinearPartition-C | 0.523 | 0.542 | 0.542 |
|  | LinearPartition-V | 0.499 | 0.453 | 0.589 |

In order to comprehensively evaluate the generalization capability of our RSSM series, we obtain two external datasets Sato and Kato (2021): the bprna-1m dataset and an Rfam-derived dataset consisting of RNA families not included in the bprna training set. Here we only give a succinct summary of our findings; see Suppl. section B and Table S2, S3 and S4 for more details on these external datasets and our experiments.

On the Rfam-derived dataset featuring new RNA families, RSSM_d64_ and RSSM_d64,T_ have limited generalization capability compared to traditional approaches. Nevertheless, comparable or even superior performance is achieved by a novel RSSM ensemble referred as RSSM_d64,mix_, which combines a diverse selection of model hypotheses that aim to more thoroughly leverage computationally predicted RNA secondary structures. Models in this mixed ensemble are either trained on bprna dataset as usual in the earlier section, or finetuned with lower learning rates and optionally augmented with sequences sampled from our curated Rfam pretraining data, with labels being predicted basepairing probabilities rather than real RNA structures.

Including these computationally predicted structures into finetuning slightly compromises the performance of RSSM_d64,mix_ on real RNA structures in the bprna-1m dataset. However, a more balanced performance can now be achieved between old and new RNA families in the bprna-1m and Rfam-new-families dataset. In fact, RSSM_d64,mix_ has caught up with or even surpassed state-of-the-art traditional approaches on short and medium RNAs in the Rfam derived dataset.

It is worth emphasizing that the improvement of RSSM_d64,mix_ is achieved without having access to any real RNA structures in newer RNA families, which indicates that our RSSMs and presumably other learning based methods can adapt to new RNA families and domains of structures even if experimentally determined RNA structures are not available for those families. This is achieved by leveraging a larger pool of RNAs with computationally predicted structures produced by LinearPartition Zhang *et al*. (2020).

We also evaluate our RSSMs on recent datasets of RNA secondary structures determined by high resolution experimental techniques, such as crystallized structures deposited in PDB database and other NMR structures. These additional results are presented in Suppl. section D, and we observe similar performance gains of RSSM_d64,mix_. Different contact map discretization techniques potentially have a large impact on the final performance, which we analyze in Suppl. section E. We find out that our RSSMs are amenable to a wide variety of discretization strategies at low probability cutoffs.

### 5.3 Performance of RSSM on long RNA sequences

The length of RNA sequences included for pretraining or finetuning is either capped at 1000 or 500 nts respectively for the sake of computational efficiency. Therefore, prediction accuracy on longer RNA sequences (up to 4381 nts) needs to be assessed. As results in Table S4 show, our most sophisticated ensemble model RSSM_d64,mix_ underperforms compared to the best performing traditional algorithms (usually LinearPartition).

Without soliciting additional supervision from these much longer sequences, we seek a simple workaround by adding LinearPartition into our ensemble model in order to leverage its built-in inductive bias on larger RNA structure decomposition. This leads to a hybrid model referred as RSSM_d64,mix,LPC_ which comprehensively surpasses RSSM_d64,mix_ on long RNA sequences in both dataset.

Furthermore, we adopt a sliding window based local structure prediction approach, considering the fact that RNA basepair spans are effectively capped during training, which gives rise to a reasonable conjecture that RSSM cannot reliably predict longer basepairs. Therefore, in the hope of trading the discovery of possible longer range interactions for more accurate local RNA structures, we average basepairing probabilities predicted within many overlapped local windows (Equation 5), as originally proposed in RNAplfold Bernhart *et al*. (2006).

Combining the sliding window approach with our RSSMs tangibly improves their performance on long RNA sequences in both bprna-1m and Rfam-new-families, as shown in Suppl. section C and Table S5.

### 5.4 Performance of alignment based models

Performance of our alignment based RSSMs is shown in Table 3, while full result is contained in Suppl. section I. Details about the role played by consensus secondary structures annotation inside the RNAcmap pipeline Zhang *et al*. (2021), which is a crucial step for obtaining MSAs, are in section Task description.

**Table 3:** Performance of our alignment based models on the bprna test set. Subscript also indicates the type of consensus secondary structure used to annotate MSAs (rnafold or dl).

| Mode | Model | Test set |  |  |
| --- | --- | --- | --- | --- |
|  |  | F1 | Precision | Recall |
| Single models | RSSM<sub>d64,rnafold</sub> | 0.733±0.003 | 0.742±0.002 | 0.742±0.004 |
|  | RSSM<sub>d64,T,rnafold</sub> | 0.740±0.004 | 0.755±0.009 | 0.745±0.005 |
|  | RSSM<sub>d64,dl</sub> | 0.742±0.001 | 0.764±0.002 | 0.738±0.002 |
|  | RSSM<sub>d64,T,dl</sub> | 0.747±0.002 | 0.769±0.002 | 0.743±0.004 |
| Ensemble models | RSSM<sub>d64,rnafold</sub> | 0.746 | 0.766 | 0.747 |
|  | RSSM<sub>d64,T,rnafold</sub> | 0.752 | 0.779 | 0.749 |
|  | RSSM<sub>d64,dl</sub> | 0.749 | 0.779 | 0.741 |
|  | RSSM<sub>d64,T,dl</sub> | 0.754 | 0.782 | 0.747 |
| End-to-end model | MSA-RSSM<sub>d64,T,rnafold</sub> | 0.777 | 0.793 | 0.776 |
| Baselines | SPOT-RNA2-IL0 | 0.727 | 0.731 | 0.723 |
|  | SPOT-RNA2-IL1 | 0.734 | 0.780 | 0.693 |
|  | SPOT-RNA2-IL2 | 0.726 | 0.716 | 0.737 |
|  | RNAalifold | 0.552 | 0.695 | 0.498 |
|  | CentroidAlifold | 0.619 | 0.611 | 0.679 |
|  | IPknot | 0.627 | 0.680 | 0.623 |

Compared to single sequence based RSSMs, the gain from simply adding covariance features extracted from MSAs is significant (see Figure 4 A), irrespective of the type of consensus secondary structure annotated within these MSAs (indicated by the subscript, e.g. *dl* stands for our own deep learning model RSSM_d64,T_). Performance gains are significant and reliable especially when the number of effective sequences in the MSA is sufficiently large (*N*_eff_ *>* 50) as shown in Figure 4 B and C. Among other contributing factors, finetuning on top of evolutionarily pretrained RSSMs and ensemble modeling have also tangibly contributed to the performance.

**Figure 4:**
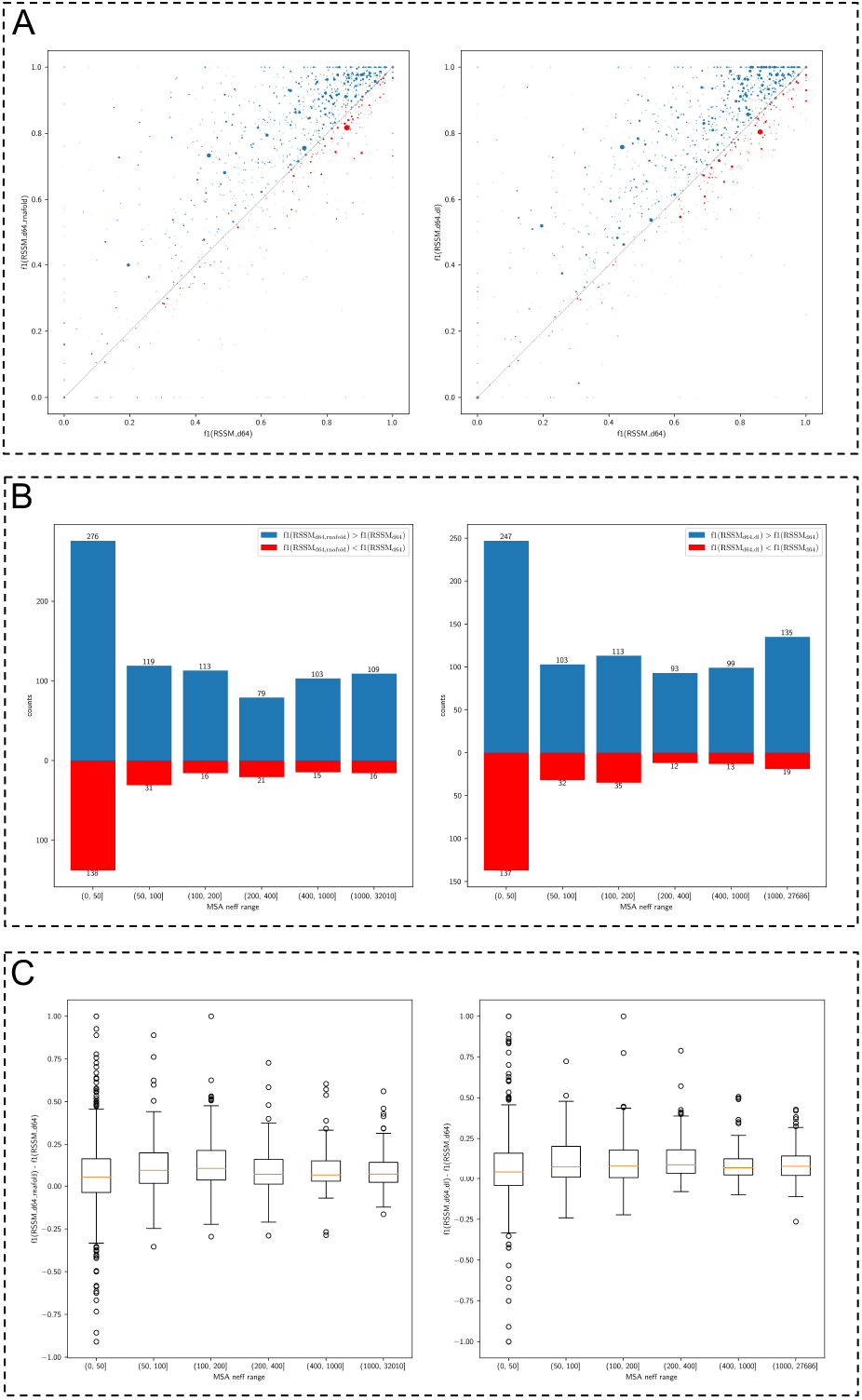
(A) Performance of alignment based RSSM_d64,rnafold_ and RSSM_d64,dl_ compared to single sequence based RSSM_d64_. Point size indicates *N*_eff_ of MSA. (B), (C) The effect of *N*_eff_ on model performance illustrated in bar plots and box plots.

The type of consensus secondary structure lays a large impact on MSAs and consequently influences the performance (see section Task description and Figure 2 for a better understanding). In particular, consensus secondary structures annotated by our own deep learning tool i.e. one RSSM_d64,T_ replicate, appear to entail more powerful MSAs, which is indicated by the performance gain of RSSM_d64,dl_ over RSSM_d64,rnafold_. In Suppl. section H and Figure S10, we directly extract co-conservation signals through DCA from MSAs as a mean to predict RNA basepairs. Once again, our results suggest the superiority of deep learning annotated MSAs.

However, further analysis on the robustness of alignment based RSSMs (Supp. section J.1) reveals that RNAfold-annotated MSAs are more suitable for training purposes, as RSSM_d64,T_ annotated MSAs can induce severe overfitting and cause instability of trained RSSMs on the RNAfold annotated MSAs (Table S8). On the other hand, RSSMs trained with RNAfold annotated MSAs are more robust and in fact, perform better on RSSM_d64,T_ annotated MSAs. We seek to better understand this phenomenon in section J.1, and point out a possible source of perturbation in SPOT-RNA2 intermediate features that can undermine its prediction accuracy.

To fully integrate evolutionary information from raw MSAs, we combine MSA transformer Rao *et al*. (2021) with RSSM which leads to an end-to-end model referred as MSA-RSSM. More details on this model and a dedicated novel ensembling strategy through repeated random MSA subsampling can be found in Suppl. section I. We pretrain the MSA transformer using Rfam MSAs as part of our pretraining strategies (section “Pretraining strategies”), before the whole MSA-RSSM is finetuned on RNAfold-annotated MSAs. Coupled with the novel ensembling strategy, MSA-RSSM produces a new state-of-the-art on the benchmark in Table 3 and S7.

Furthermore, we investigate the robustness of our RSSMs on shallow MSAs (see Suppl. section J.2 and Figure S11). Both RSSM_d64,T,rnafold_ and MSA-RSSM_d64,T,rnafold_ are relatively robust to shallow MSAs as they outperform single sequence RSSM_d64,T_ as long as a minimal amount of *N*_eff_ = 5 homologous sequences are retained for each MSA under the low setting, or *N*_eff_ = 10 sequences in the high setting. In addition, MSA-RSSM_d64,T,rnafold_ has been shown to better utilize evolutionary information than the covariance feature based RSSM.

### 5.5 Generalization of alignment based RSSMs

We also evaluate our alignment based RSSMs on new RNA families, similar to single sequence based models in B, except we focus on Rfam-derived dataset and only consider medium to long RNA sequences. Full results are presented in Suppl. section K, Table S9 and Table S10.

Using RNAfold or RSSM_d64,mix_ annotated MSAs, on medium RNA sequences and long RNA sequences with pseudoknots, our RSSM_d64,rnafold_, RSSM_d64,T,rnafold_ and MSA-RSSM_d64,T,rnafold_ significantly outperform all single sequence as well as alignment based baselines such as traditional RNA folding algorithms and DCA. In particular, our end-to-end MSA-RSSM_d64,T,rnafold_ performs the best in the benchmark on medium RNA sequences between 151 and 500 nts, and its efficacy is most impressive on RNAs with pseudoknots where we observe consistent performance gain over the best covariance based RSSM, and 11% over the best single sequence model.

Two examples of such performance gain is shown in Figure 5, featuring less well-studied RNAs from two novel RNA families in the Rfam derived dataset Sato and Kato (2021). The observed benefits of leveraging evolutionary covariance features and hybrid MSA-RSSM architecture include the reduction of false positive predictions (panel A), and identification of longer range basepairing interactions as well as pseudoknots (panel B).

**Figure 5:**
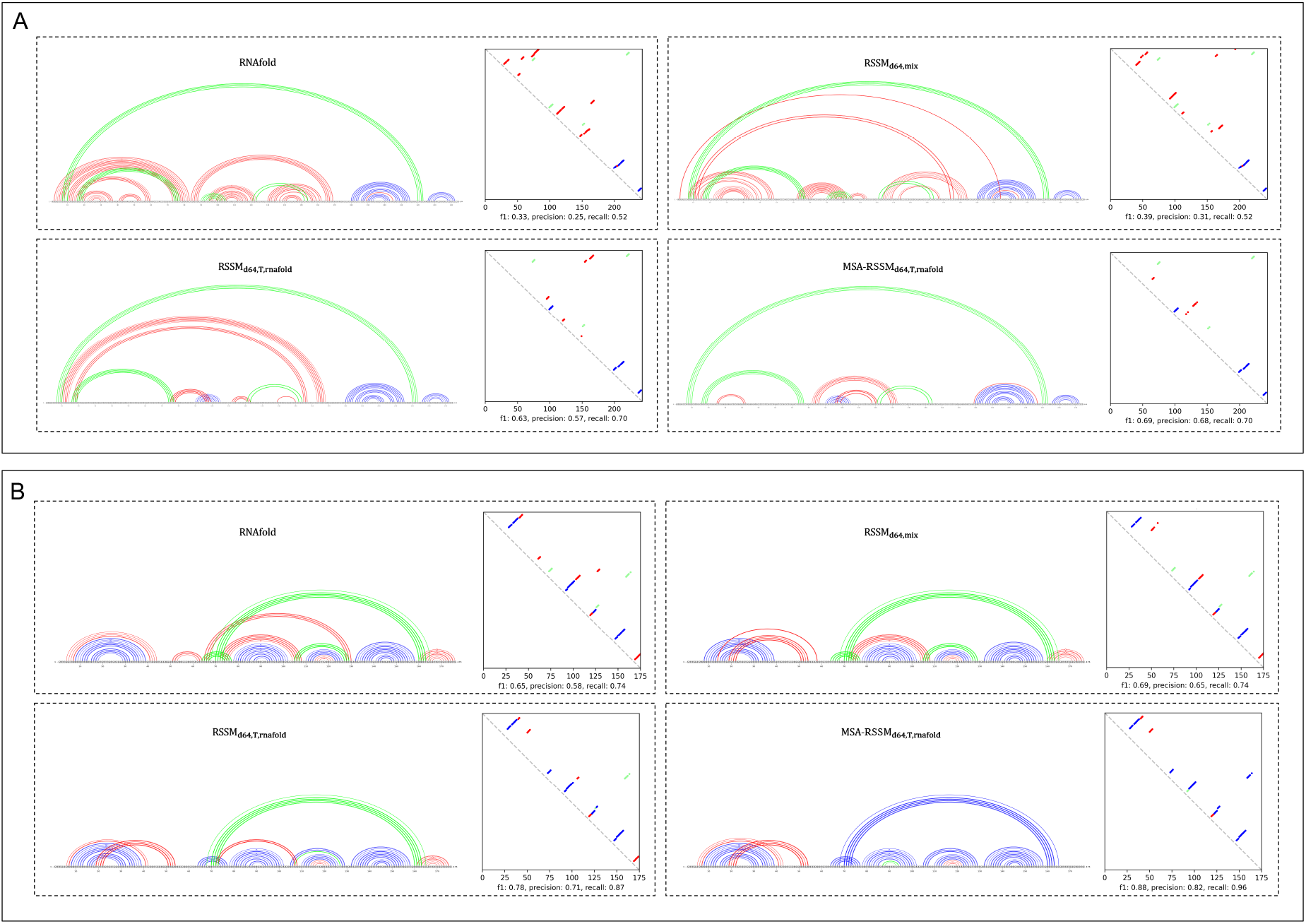
Leveraging evolutionary information improves prediction accuracy of many RNA secondary structures in the Rfam derived dataset. Panel A: an example of a small bacterial non-coding RNA (Rfamid: RF02673, seqid: ABYA01000314.1/2329-2086), with 330 non-redundant homologs in the MSA. Panel B: another example with pseudoknots and 16 non-redundant homologs in the MSA (Rfamid: RF03060, seqid: URS0000D692E3.12908/1-176). Note that MSA-RSSM_d64,T,rnafold_ correctly identifies pseudoknots. Color scheme: blue dots/lines indicate true positives; red dots/lines indicate false positives; green dots/lines indicate false negatives. RNA structures are plotted by VARNA software Darty *et al*. (2009).

Overall, comparing alignment based to single sequence based RSSMs reveals that the leveraging evolutionary information substantially improves performance on new RNA families. Our alignment based RSSMs are also more effective at incorporating evolutionary information than traditional methods.

## 6 Discussion

With evolutionary information integrated pretraining, our deep learning models obtain state-of-the-art performance across a range of benchmark tasks that span single sequence and alignment based structure prediction, and possess superior generalization capability rivaling or surpassing the state-of-the-art traditional RNA folding algorithms in new RNA families. Nevertheless, we discuss the limitation of our RSSMs and point out some important future research directions.

First, despite the superior predictive power demonstrated by our single sequence based RSSMs, on small and medium RNA secondary structures, and especially those with pseudoknots and non-canonical basepairing interactions, our approaches still lack the capability of handling much longer RNAs, emitting largely comparable performance with the state of the art thermodynamic models in the long sequence regime. Although the sliding window method described in section “Performance of RSSM on long RNA sequences” and Suppl. section C consistently improves RSSMs prediction for RNA sequences longer than 2000 nts, it sacrifices the model’s ability of identifying potential longer range interactions, instead trading it for more accurate local structure characterizations.

Based on these observations, we believe more impactful future research should focus on scaling computational approaches to longer RNA sequences beyond 1000 nts, possibly by integrating more biologically inspired inductive bias similar to MXFold2 (Sato *et al*., 2021), although the computational overhead incurred by such integration would become a major concern. The special accommodation of pseudoknots, due to the nature of structural decomposition under the nearest neighbour framework, has yet to be resolved as well.

Improvements to our proposed pretraining strategies can also be conceived on the single sequence level as well as alignment level. In particular, our analysis in section “Performance of alignment based models” and Suppl. section J.1 reveal that MSAs with weaker CSS annotation lead to more robust models. This suggests that our alignment based RSSMs can potentially benefit from larger scale pretraining beyond the resources available in the current Rfam database, possibly by leveraging MSAs obtained directly from blastn without any CSS annotations, similar to the type of MSAs employed in protein structure related research (Jumper *et al*., 2021; Rao *et al*., 2021). We point out that only using blastn also reduces the computation demand compared to the full RNAcmap pipeline.

Finally, in Suppl. section L we briefly showcase the detection of spurious homologs through our pretrained MSA-transformer. Future investigation down this route on the removal of such detected spurious homologs as a layer of quality control may lead to further improvement in our alignment based model RSSMs.

We hope our work opens the door to future research on computational RNA secondary structure prediction, as more RNA structure data can be made available to support deep learning methods, accompanied with dedicated modelling techniques and pretraining strategies for single sequence as well as alignment based models to better tackle the challenges posed by RNAs from distant families and the much longer sequence regime.

## Acknowledgement

We thank Compute Canada (Narval, Beluga, Graham and Cedar) for providing the computational resources. We also thank Jérôme Waldispühl for the discussion.

## Appendix

### A A Training details

#### A.1 A.1 Supervision on intermediate feature maps

In addition to the final RNA contact map representation, RSSM generates a set of downsampled feature maps which have condensed information about potential long range basepairs within or between large patches of merged contact map areas. These features are eventually merged into the final RNA contact map representation through the horizontal shortcut and expansive path. Nevertheless, we provide auxiliary supervision to these intermediate feature maps in the hope of better extracting and integrating these information to improve the final RNA contact map prediction. Our auxiliary targets are in the form of basepairing intensity scores that can be obtained from the ground truth binary RNA contact map, as illustrated in Figure 3 B. The loss function can be expressed as follow:

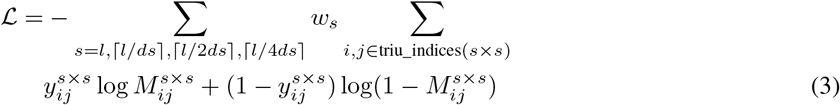

where *s* denotes the spatial dimensionality of the full RNA contact map or downsampled feature map. and *w*_*s*_ are weights to balance the loss from different maps. When the map is in its full resolution, i.e. 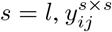 are binary labels, with 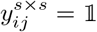 (i forms a basepair with j). In its downsampled versions, for example 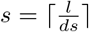,

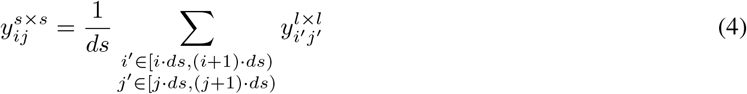

considering that at most *ds* basepairs could be formed between two stretches of distant RNA segments of length *ds*, thereby rendering 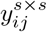 a measurement of basepairs concentration within a merged patch in the ground truth RNA contact map.

*M*^*s*×*s*^ contains basepairing probabilities or intensities predicted from the corresponding RNA contact map representation or downsampled feature maps. Each *M*^*s*×*s*^ employs a separate classification head implemented as a three-layer multilayer perceptron (MLP), followed by sigmoid activation. Each fully-connected layer in the MLP is preceded by layer normalization, nonlinear activation and dropout.

**Figure S1:**
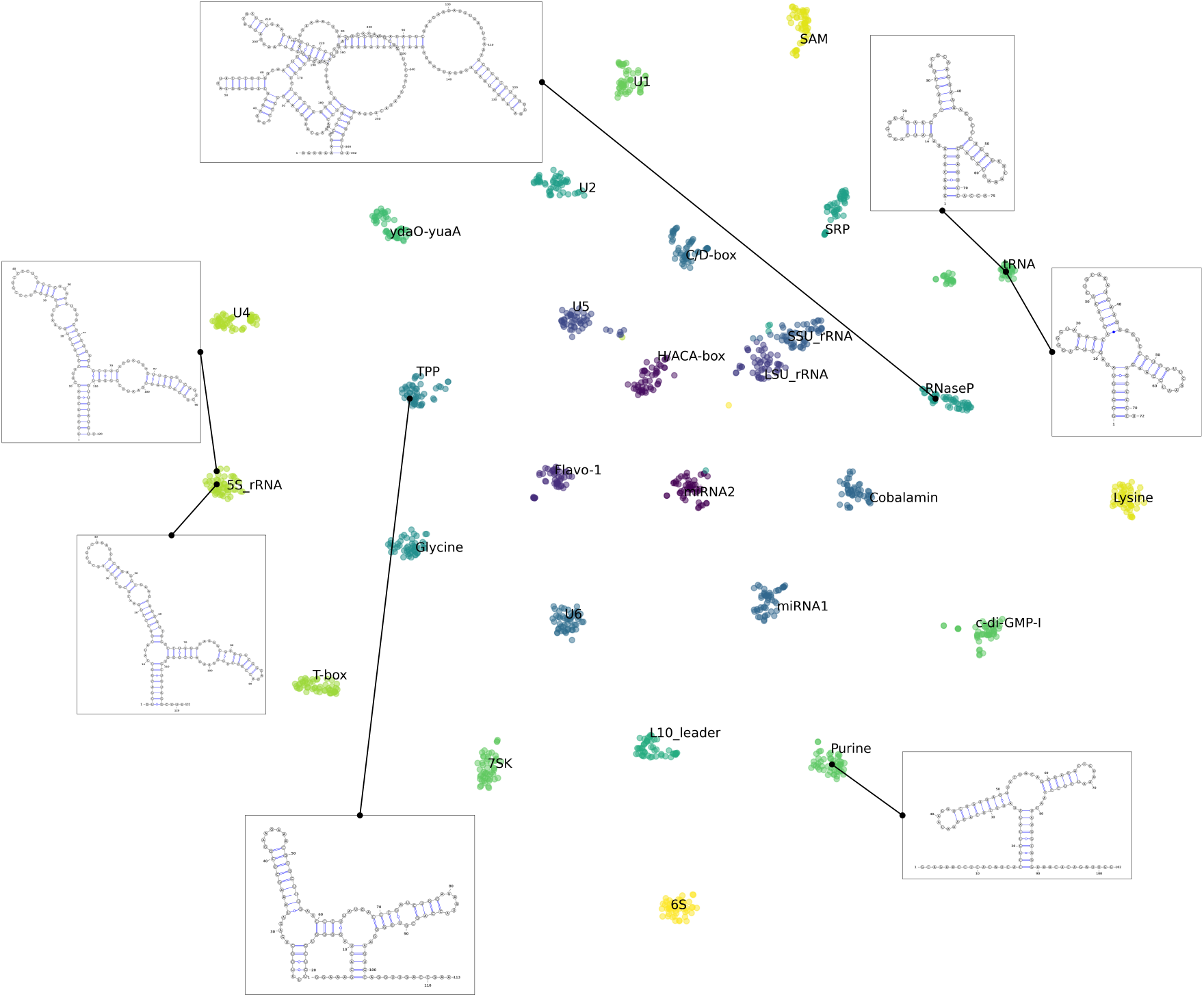
We project the RNA embeddings learnt through single sequence pretraining to two dimensional space via t-SNE, a dimensionality reduction algorithm. These projected embeddings are originally obtained from the held-out fraction of our pretraining data. Clusters of distinct RNA families can be clearly identified, as a result of our non-coding RNA classification objective which is part of the pretraining strategy. RNA secondary structures are also shown for a number of representative RNA families that include tRNA, RNase P, Purine, TPP riboswitch, and small 5s ribosomal RNAs. These structures are predicted by our own RSSM model (RSSM_d64,mix_, which is introduced in section “Generalization to new RNA families” and section B), and drawn by VARNA.

#### A.2 A.2 Hyperparameters and training configurations

**Table S1:**
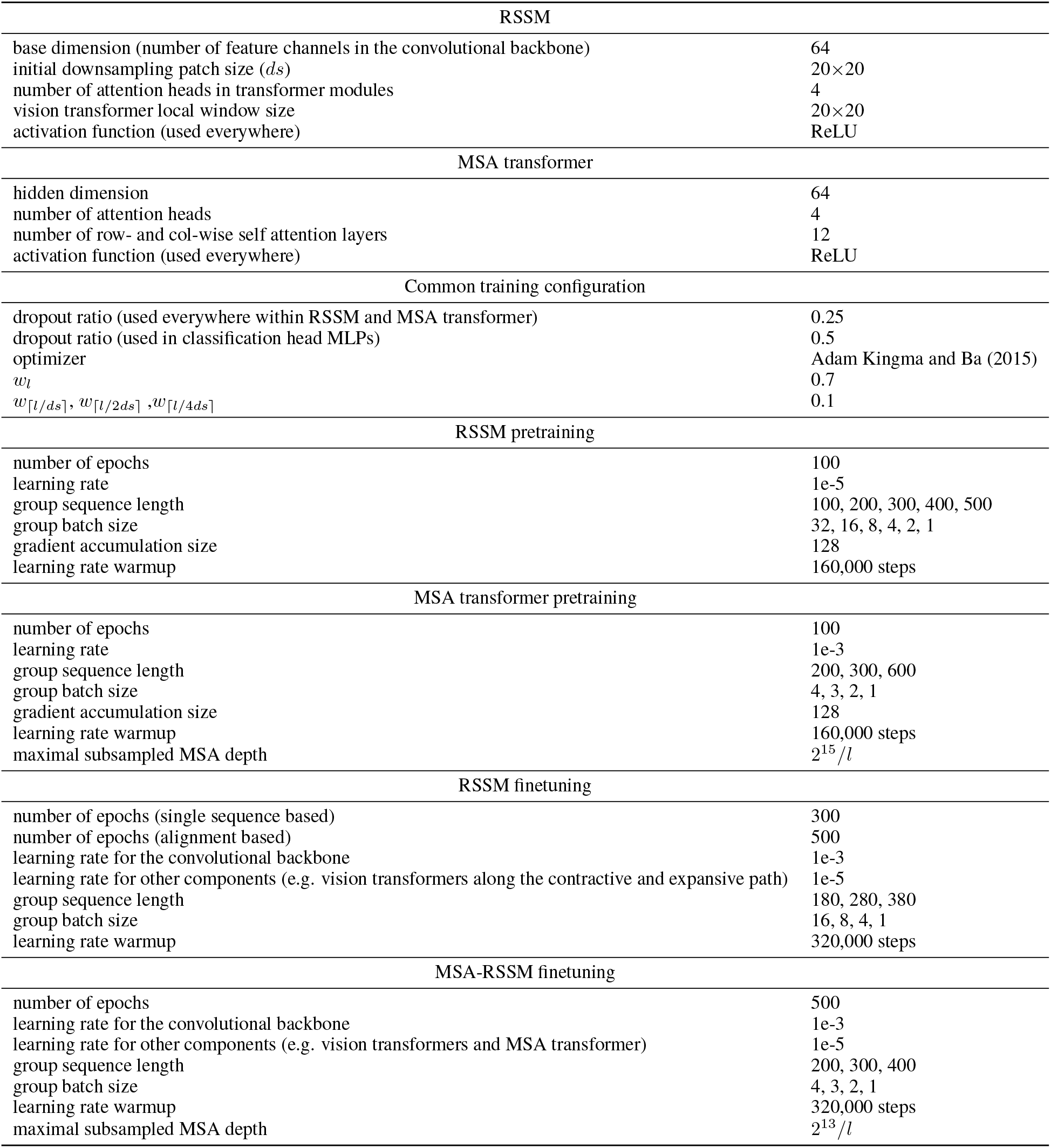
Below are hyperparameters and training configurations for RSSM and MSA transformer. For the sake of training efficiency, we group RNA sequences into different ranges of length indicated in the “group sequence length” entry. For example, “180, 280, 380” indicates four length ranges that are (0, 180], (180, 280], (280, 380] and (380, 500]. Each length group has a different batch size indicated by “group batch size”.

### B B RSSM generalization performance in external dataset

**Table S2:**
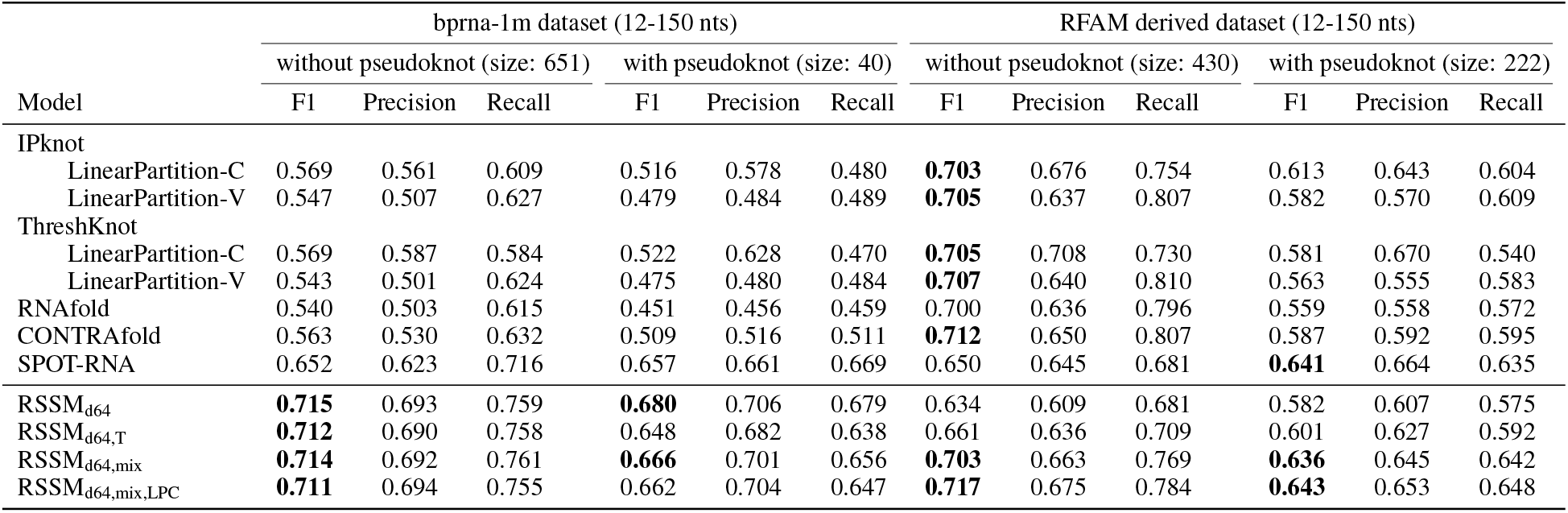
Performance of RSSMs and baselines including SPOT-RNA and a variety of traditional RNA folding algorithms in bprna-1m dataset and Rfam derived dataset, featuring short RNA sequences between 12 to 150 nts with or without pseudoknots.

We measure the generalization capability of our RSSMs on two recent dataset curated by Sato and Kato, 2021.

- Bprna-1m dataset is extracted from the same online database Danaee *et al*. (2018) that was used earlier in Singh *et al*., 2019 to compile bprna training, validation and test sets. Nevertheless, having removed RNA sequences that share more than 80% similarity to those in the existing bprna dataset, the newer bprna-1m dataset can be used to measure the performance of our RSSMs on new RNA sequences from the existing RNA families.
- Rfam derived dataset is curated from the latest Rfam database (version 14.5) using consensus secondary structures as ground truth, and it only contains sequences from new RNA families that are absent in the bprna dataset. Therefore, Rfam derived dataset is employed to evaluate the generalization capability of our RSSMs in new RNA families, and to see if a balanced performance can be achieved between existing and new RNA families.

In addition to the data curation by Sato and Kato, 2021, we further control sequence similarity in bprna-1m and Rfam derived dataset and remove RNA sequences therein that share at least 80% similarity against our own pretraining data, using cd-hit-est-2d.

We follow the same evaluation scheme in Sato and Kato, 2021 and compare our RSSMs to baselines including SPOT-RNA and a variety of RNA folding algorithms. RNAs in these two dataset are grouped into different types according to their length and the existence of pseudoknots, as shown in Table S2, Table S3 and Table S4. We include fours types of RSSM into the comparison, which are RSSM_d64_, RSSM_d64,T_, RSSM_d64,mix_ and RSSM_d64,mix,LPC_. Each one is a five-model ensemble:

- RSSM_d64_ is composed by directly trained models using bprna training set (as in in Table 2).
- RSSM_d64,T_ is composed by finetuned models following pretraining (as in in Table 2).
- RSSM_d64,mix_ is a mixture of directly trained and finetuned models, with its formula carefully chosen to balance its generalization performance in existing and new RNA families. We will elucidate its exact composition later in greater details.
- RSSM_d64,mix,LPC_ is essentially the same RSSM_d64,mix_, but with additional LinearPartition-V basepairing probabilities incorporated into our ensemble model as a crude form of leveraging biological priors.

For each RNA type in Table S2, Table S3 and Table S4, the best performing method or those within 2% difference of the best are marked with boldface. For the simplicity of analysis, we compare their performance by the F1 score. Across all RSSM series, a threshold of 0.3 is used for all RNA types in the bprna-1m dataset, and a threshold of 0.2 is used for all RNA types in the Rfam derived dataset.

**Table S3:** Performance of RSSMs and baselines including SPOT-RNA and a variety of traditional RNA folding algorithms on medium RNA sequences between 151 to 500 nts with or without pseudoknots in bprna-1m dataset and Rfam derived dataset.

| Model | bprna-1m dataset (151-500 nts) |  |  |  |  |  | RFAM derived dataset (151-500 nts) |  |  |  |  |  |
| --- | --- | --- | --- | --- | --- | --- | --- | --- | --- | --- | --- | --- |
|  | without pseudoknot (size: 202) |  |  | with pseudoknot (size: 72) |  |  | without pseudoknot (size: 169) |  |  | with pseudoknot (size: 70) |  |  |
|  | F1 | Precision | Recall | F1 | Precision | Recall | F1 | Precision | Recall | F1 | Precision | Recall |
| IPknot |  |  |  |  |  |  |  |  |  |  |  |  |
| LinearPartition-C | 0.451 | 0.445 | 0.486 | 0.384 | 0.404 | 0.375 | 0.506 | 0.453 | 0.610 | 0.503 | 0.475 | 0.549 |
| LinearPartition-V | 0.424 | 0.382 | 0.503 | 0.413 | 0.386 | 0.457 | 0.493 | 0.404 | 0.667 | 0.456 | 0.393 | 0.555 |
| ThreshKnot |  |  |  |  |  |  |  |  |  |  |  |  |
| LinearPartition-C | 0.455 | 0.490 | 0.461 | 0.381 | 0.448 | 0.343 | **0.512** | 0.494 | 0.572 | 0.490 | 0.503 | 0.500 |
| LinearPartition-V | 0.426 | 0.383 | 0.504 | 0.407 | 0.384 | 0.444 | 0.498 | 0.408 | 0.670 | 0.453 | 0.392 | 0.546 |
| RNAfold | 0.415 | 0.373 | 0.493 | 0.391 | 0.367 | 0.429 | 0.486 | 0.400 | 0.651 | 0.433 | 0.377 | 0.520 |
| CONTRAFold | 0.445 | 0.414 | 0.510 | 0.419 | 0.406 | 0.441 | 0.510 | 0.428 | 0.664 | 0.473 | 0.420 | 0.556 |
| SPOT-RNA | 0.448 | 0.476 | 0.456 | 0.455 | 0.530 | 0.409 | 0.439 | 0.443 | 0.462 | 0.446 | 0.482 | 0.427 |
| RSSM<sub>d64</sub> | 0.535 | 0.542 | 0.554 | 0.600 | 0.625 | 0.590 | 0.437 | 0.394 | 0.515 | 0.493 | 0.476 | 0.524 |
| RSSM<sub>d64,T</sub> | **0.568** | 0.561 | 0.603 | **0.627** | 0.634 | 0.633 | 0.475 | 0.415 | 0.584 | 0.480 | 0.456 | 0.521 |
| RSSM<sub>d64,mix</sub> | 0.551 | 0.558 | 0.574 | 0.612 | 0.628 | 0.608 | 0.505 | 0.437 | 0.627 | **0.510** | 0.480 | 0.555 |
| RSSM<sub>d64,mix,LPC</sub> | 0.548 | 0.558 | 0.571 | 0.608 | 0.629 | 0.600 | **0.521** | 0.452 | 0.644 | **0.514** | 0.483 | 0.560 |

#### B.1 B.1 RSSM_d64_ performs well on sequences shorter than 500 nts in the bprna-1m dataset, but generalize poorly on longer RNA sequences and new RNA families in the Rfam derived dataset

Compared to a wide selection of baselines, RSSM_d64_ always emits better or comparable performance on sequences below 500 nts in the bprna-1m dataset, which represent the group of RNA sequences similar to its training set. On the other hand, in Rfam derived dataset and on longer RNAs, RSSM_d64_ exhibits extremely limited accuracy, especially compared to RNAfold which is a traditional thermodynamic model that is built on the decomposition of nested RNA secondary structures. RSSM_d64_ also falls short of CONTRAfold which similarly encodes strong inductive bias of RNA secondary structures through a generalized form of stochastic context free grammar. Both approaches rely on strong biological priors which cannot fully explain observed RNA secondary structure data, but they do provide a set of biologically well-grounded principles for RNA folding which can be extended to any new RNA families.

IPknot and ThreshKnot have also demonstrated superior performance over RSSM_d64_, and these approaches can be understood as discretization techniques on top of basepairing probabilities in a predicted RNA contact map. We only apply IPknot and ThreshKnot on basepairing probabilities predicted by LinearPartition, which usually leads to better performance than other alternatives such as McCaskill’s algorithm or CONTRAfold’s native scoring system Sato and Kato (2021).

Compared to SPOT-RNA, our RSSM_d64_ has shown better performance on medium and long RNAs with pseudoknots in the Rfam derived dataset, while maintaining largely identical performance elsewhere, with the exception of short pseudoknotted structures.

In general, models with biological priors have consistently outperformed deep learning selections that are SPOT-RNA and RSSM_d64_. This indicates that given limited amount of training data, existing deep learning approaches that are trained from scratch would not outperform traditional approaches in new RNA families or on longer sequences that are absent in the training set, unless more data is made available to properly represent these newer domains of RNA structures.

#### B.2 B.2 Transfer learning improves generalization capability across all dataset and RNA types

Adding RSSM_d64,T_ to the comparison, we observe that performance is improved in almost every setting, especially for long RNA sequences and new RNA families in the Rfam derived dataset. In particular, RSSM_d64,T_ outperforms its direct learning counterpart RSSM_d64_ as well as the deep learning baseline SPOT-RNA on almost every RNA type in new RNA families, with the exception of short pseudoknotted sequences.

Nevertheless, the gap between RSSM_d64,T_ and the variety of traditional RNA folding algorithms still exists. In an attempt to fully tap into the power of pretraining, and for the purpose of balancing the performance of our model between old and new RNA families, between shorter and longer RNAs, we compose a novel RSSM ensemble model with experts addressing different aspects of these generalization tasks, as opposed to having a simple ensemble of replicate models.

The idea of composing an ensemble with different hypotheses stems from a crucial observation on SPOT-RNA, which is originally an ensemble of five different deep learning architectures. While the whole SPOT-RNA ensemble has shown an impressive performance on the these generalization tasks, we find out that each single model performs much worse, which seems to suggest that ensembling diverse model hypotheses may lead to a more dramatic increase in the overall performance.

Therefore, we propose to build an ensemble with mixed hypotheses referred as RSSM_d64,mix_. While we make no attempt at searching for new model architectures, we simply train our basic RSSM with different learning schedules. Our RSSM_d64,mix_ is an ensemble of the following models, all trained to 300 epochs:

- One directly trained model from RSSM_d64_.
- One finetuned model from RSSM_d64,T_.
- Two finetuned models with a smaller base learning rate at 1e-5, in order to help RSSM to retain more knowledge of the diverse RNA secondary structures it has seen during pretraining.
- One finetuned model but with subsampled Rfam pretraining data directly complementing each epoch.

The mixture in our RSSM_d64,mix_ is selected with the purpose of balancing the information beneficial to the existing RNA families in bprna dataset, or to the newer RNA families in Rfam derived dataset. In particular, finetuned models with lower learning rates tend to retain more information from its pretraining, while complementing the finetuning process directly with subsampled Rfam pretraining data also boosts the generalization capability of our RSSMs on other RNA sequences in newer RNA families.

#### B.3 B.3 Ensembling diverse hypotheses improves generalization in new RNA families and helps to obtain more balanced performance between existing and new RNA families

Our mixed ensemble RSSM_d64,mix_ is able to maintain a more balanced performance between bprna-1m and Rfam derived dataset. In particular, improvement in Rfam derived dataset is the most substantial, with RSSM_d64,mix_ outperforming all aforementioned deep learning selections across all RNA types by a large margin. Compared to the state-of-the-art traditional RNA folding algorithms, RSSM_d64,mix_ demonstrates superior performance on short and medium pseudoknotted RNA sequences, which can be understood from the fact that RSSM naturally supports pseudoknot predictions, while RNAfold, CONTRAfold and LinearPartition cannot.

On the other hand, while RSSM_d64,mix_ emits largely comparable or slightly lower performance on short and medium RNA sequences without pseudoknots, it falls short on longer pseudoknotted RNA sequences not only in the Rfam derived dataset but also in the bprna-1m dataset, which is not unexpected given the fact that the maximal length of RNA sequences included throughout pretraining or finetuning is capped at 1000 or 500 nts respectively.

In bprna-1m dataset, RSSM_d64,mix_ has largely comparable or slightly lower accuracy compared to the deep learning selections RSSM_d64_ or RSSM_d64,T_. Since our Rfam pretraining data only contain predicted basepairing probabilities that cannot reflect pseudoknots or non-canonical basepairs in actual RNA secondary structures, introducing these computationally predicted structures into finetuning has slightly compromised the performance of RSSM_d64,mix_ on real RNA secondary structures in the bprna and bprna-1m dataset. Nevertheless, RSSM_d64,mix_ is still able to outperform all traditional RNA folding algorithms in both datasets.

Overall, given the balanced performance of RSSM_d64,mix_ in the more challenging Rfam derived dataset, reaching or surpassing the state-of-the-art traditional RNA folding algorithms on short and medium RNA sequences, in addition to the fact that RSSM_d64,mix_ has outperformed all other deep learning models on long RNA sequences by a large margin, it is fair to say RSSM_d64,mix_ is the most useful RNA secondary structure prediction tool we have built so far.

#### B.4 B.4 Improving the prediction for longer RNAs by incorporating LinearPartition basepairing probabilities

Throughout Table S4, our RSSM_d64,mix_ has fallen short on longer RNA sequences compared to traditional RNA folding algorithms. While adding biological priors to our RSSM in the form of RNA secondary structure decomposition rules represents a promising direction for improvement, here we propose a simple workaround to add LinearPartition predicted basepairing probabilities to our RSSM_d64,mix_, considering the benevolent effect of model ensemble averaging which is discussed in section F.

We refer to this LinearPartition augmented approach as RSSM_d64,mix,LPC_ and it has a systemically improved generalization capability in Rfam derived dataset, obtaining comparable or state-of-the-art performance across all RNA types compared to the traditional RNA folding algorithms, with the exception of long nonpseudoknotted RNA sequences which however only contains 1 example. The improvement is particularly impressive for longer RNA sequences, for which RSSM_d64,mix,LPC_ has obtained comparable or only slightly lower performance in bprna-1m dataset as well as Rfam derived dataset.

**Table S4:** Performance of RSSMs and baselines including SPOT-RNA and a variety of traditional RNA folding algorithms on long RNA sequences from 501 to a maximum of 4381 nts with or without pseudoknots in bprna-1m dataset and Rfam derived dataset. Note, we cannot evaluate IPknot in bprna-1m dataset due to its excessively long run time.

| Model | bprna-1m dataset (501-4381 nts) |  |  |  |  |  | RFAM derived dataset (501-917 nts) |  |  |  |  |  |
| --- | --- | --- | --- | --- | --- | --- | --- | --- | --- | --- | --- | --- |
|  | without pseudoknot (size: 155) |  |  | with pseudoknot (size: 141) |  |  | without pseudoknot (size: 1) |  |  | with pseudoknot (size: 70) |  |  |
|  | F1 | Precision | Recall | F1 | Precision | Recall | F1 | Precision | Recall | F1 | Precision | Recall |
| IPknot |  |  |  |  |  |  |  |  |  |  |  |  |
| LinearPartition-C | - | - | - | - | - | - | 0.043 | 0.042 | 0.043 | **0.450** | 0.443 | 0.468 |
| LinearPartition-V | - | - | - | - | - | - | 0.060 | 0.052 | 0.072 | 0.381 | 0.317 | 0.480 |
| ThreshKnot |  |  |  |  |  |  |  |  |  |  |  |  |
| LinearPartition-C | **0.413** | 0.429 | 0.413 | 0.404 | 0.451 | 0.405 | 0.046 | 0.049 | 0.043 | **0.444** | 0.485 | 0.422 |
| LinearPartition-V | 0.354 | 0.310 | 0.434 | 0.340 | 0.299 | 0.435 | 0.060 | 0.055 | 0.067 | 0.381 | 0.318 | 0.477 |
| RNAfold | 0.340 | 0.297 | 0.420 | 0.316 | 0.275 | 0.410 | 0.050 | 0.044 | 0.057 | 0.357 | 0.297 | 0.449 |
| CONTRAFold | 0.394 | 0.352 | 0.468 | 0.382 | 0.350 | 0.463 | 0.058 | 0.055 | 0.062 | 0.411 | 0.352 | 0.497 |
| SPOT-RNA | 0.256 | 0.273 | 0.254 | 0.280 | 0.312 | 0.282 | 0.006 | 0.008 | 0.005 | 0.346 | 0.367 | 0.330 |
| RSSM<sub>d64</sub> | 0.220 | 0.323 | 0.182 | 0.283 | 0.361 | 0.261 | 0.017 | 0.020 | 0.014 | 0.361 | 0.335 | 0.339 |
| RSSM<sub>d64,T</sub> | 0.323 | 0.330 | 0.333 | 0.368 | 0.377 | 0.396 | 0.035 | 0.036 | 0.033 | 0.402 | 0.365 | 0.452 |
| RSSM<sub>d64,mix</sub> | 0.356 | 0.415 | 0.330 | 0.392 | 0.446 | 0.388 | **0.062** | 0.067 | 0.057 | 0.421 | 0.378 | 0.479 |
| RSSM<sub>d64,mix,LPC</sub> | 0.389 | 0.454 | 0.356 | **0.413** | 0.476 | 0.402 | 0.051 | 0.054 | 0.048 | 0.432 | 0.390 | 0.485 |

Combining LinearPartition with RSSM in fact slightly reduces its performance on short and medium RNA sequences in bprna-1m dataset, possibly due to the fact that bprna-1m dataset contains a high proportion of non-canonical basepairs which cannot be predicted by LinearPartition. This leads us to another important discussion later which is to highlight the differences between bprna-1m dataset and Rfam derived dataset, and their impact on traditional RNA folding algorithms and deep learning approaches.

We point out that the selection of threshold can generate a large impact on methods such as IPknot and ThreshKnot which do not have control on the input RNA contact map basepairing probabilities. An advantage of IPknot lies in its automatic threshold selection. However, it can also consume a dramatically larger amount of computation for RNA sequences with more than 2000 nucleotides, taking up more than 250 CPU hours on a single RNA sequence and still cannot finish running. Therefore, we leave its results empty in Table S4 ^3^.

#### B.5 B.5 Difference between bprna-1m and Rfam derived datasets

The most remarkable difference between bprna-1m and Rfam derived dataset lies in their proportions of non-canonical basepairs, as shown in Figure S2. Bprna and bprna-1m datasets consistently feature up to 10% of non-canonical interactions, whereas Rfam derived dataset exclusively contains Watson-Crick and Wobble basepairs.

Another important observation can be made by noting the performance difference between RNAfold in these two dataset. Since RNAfold is a traditional energy based model with experimentally measured thermodynamic parameters, it is rather unexpected to observe such a dramatic performance gap between RNA families in bprna-1m and Rfam derived dataset. In particular, from Rfam derived dataset to bprna-1m dataset, the performance of RNAfold consistently drops more than 20% on short and medium non-pseudoknotted RNA sequences. Similar observations can be made to CONTRAfold and two LinearPartition supported methods, with respect to all RNA types. In fact, none of these methods support non-canonical interactions. It should clarified, however, CONTRAfold is by design capable of accommodating non-canonical interactions but its default mode of prediction does not support non-canonical basepairs, otherwise its overall accuracy would suffer ^4^.

Overall, these observations seem to demonstrate that non-canonical interactions pose a major challenge to existing traditional RNA folding algorithms, which may suffice to explain their large performance gap between bprna-1m and Rfam derived dataset. Deep learning approaches such as SPOT-RNA and RSSM series, in comparison, naturally support all types of basepairing interactions and such behaviour may be unfairly penalized in Rfam derived dataset. Considering that RNAs ultimately fold into tertiary structures that fulfill their biological functions in the three dimensional cellular space, RNA secondary structures as mere intermediate representation without modeling these crucial non-canonical interactions may not be able to fully capture their underlying biological properties.

**Figure S2:**
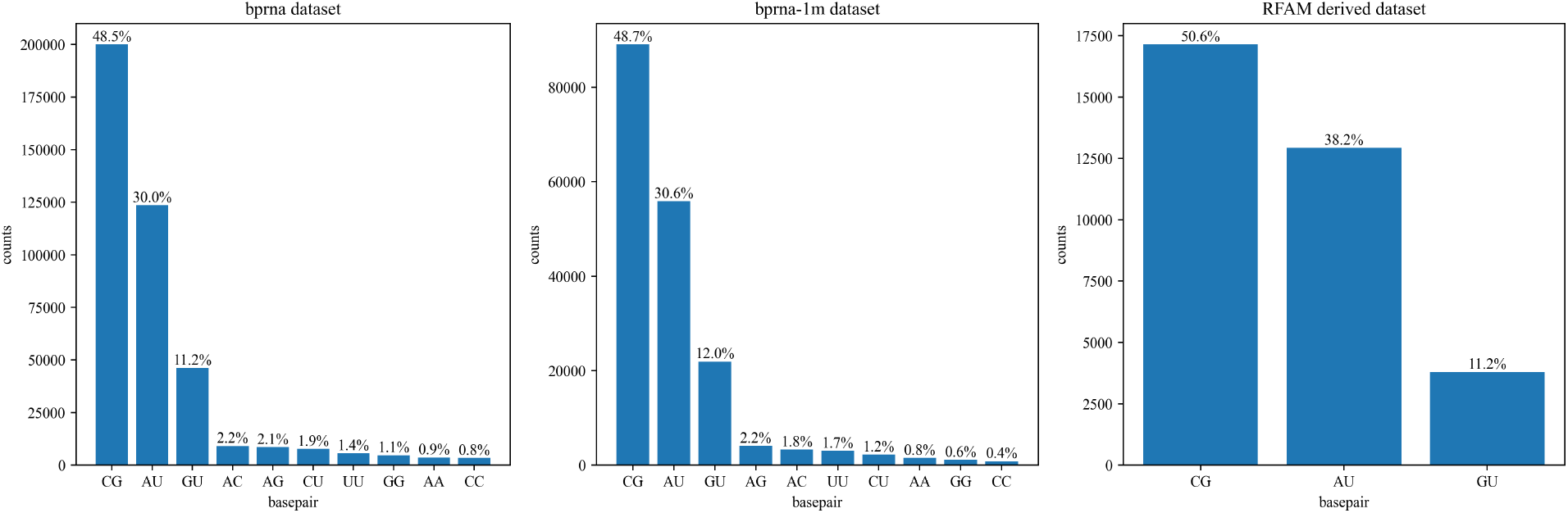
RNA secondary structures in bprna and bprna-1m dataset contain a high proportion of non-canonical basepairs which are up to 10%. Rfam derived dataset, one the other hand, does not contain any non-canonical basepairs at all.

### C C Sliding window based RNA secondary structure prediction for long RNA sequences

Consider an arbitrary basepair indexed by (*i, j*) and a window size of *L* + 1. Assuming each time we slide the window to the 3’ end by an offset of 1, then the basepairing probability averaged within all encompassed local windows can be expressed by:

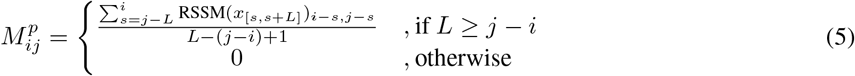

while in practice for the sake of efficiency we use larger offsets. Our results are shown in Table S5.

**Table S5:**
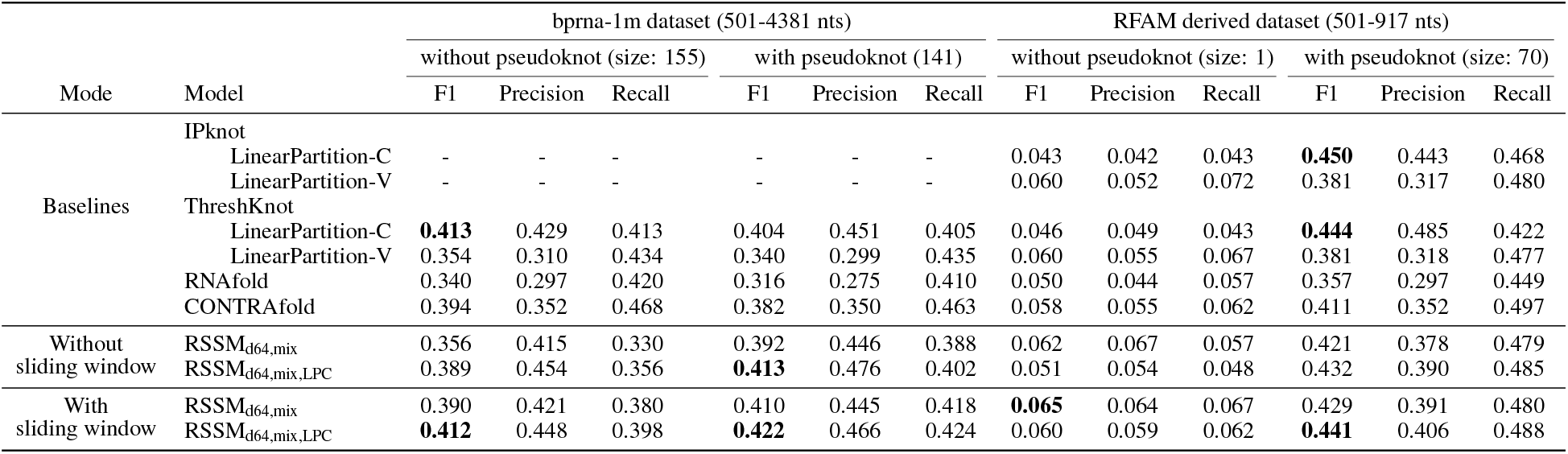
For bprna-1m dataset, we used a local window of size 1000 and a step size of 500. For Rfam derived dataset, we used a local window of size 500 and a step size of 200. Note, we cannot evaluate IPknot in bprna-1m dataset due to its excessively long run time.

We further select all 148 RNA sequences longer than 2000 nts, with or without pseudoknots, from the bprna-1m dataset, and illustrate in Figure S3 the modest but consistent improvement in the long sequence domain by using the simple yet effective sliding window approach.

**Figure S3:**
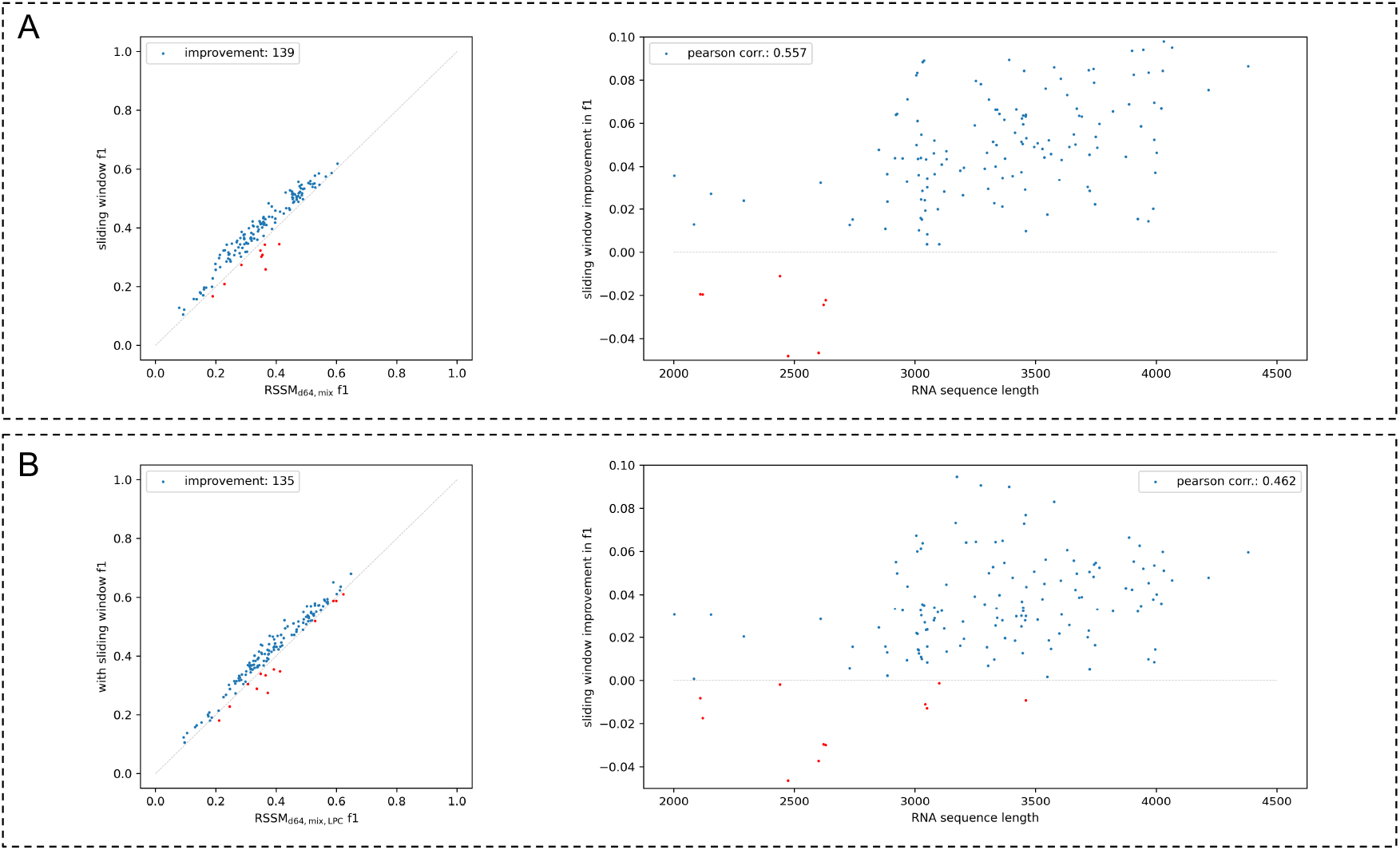
Relative f1 improvements from the sliding window approach on 148 RNAs longer than 2000 nts, with or without pseudoknots, in the bprna-1m dataset.

Figure S4 demonstrates the rationale behind the sliding window approach, which is to avoid making spurious longer range RNA basepair predictions beyond 1000 nts, and focus more on local structures hence leading to tangible improvements in RNA basepair recalls. Despite having this benevolent trade-off, our RSSMs have shown to lack the ability of correctly inferring much longer range RNA basepairing interactions, which is the case for other baselines as well as shown in Table S5. This indicates an important future direction which is to scale the current computational methods to handle much longer range RNA basepair interactions.

**Figure S4:**
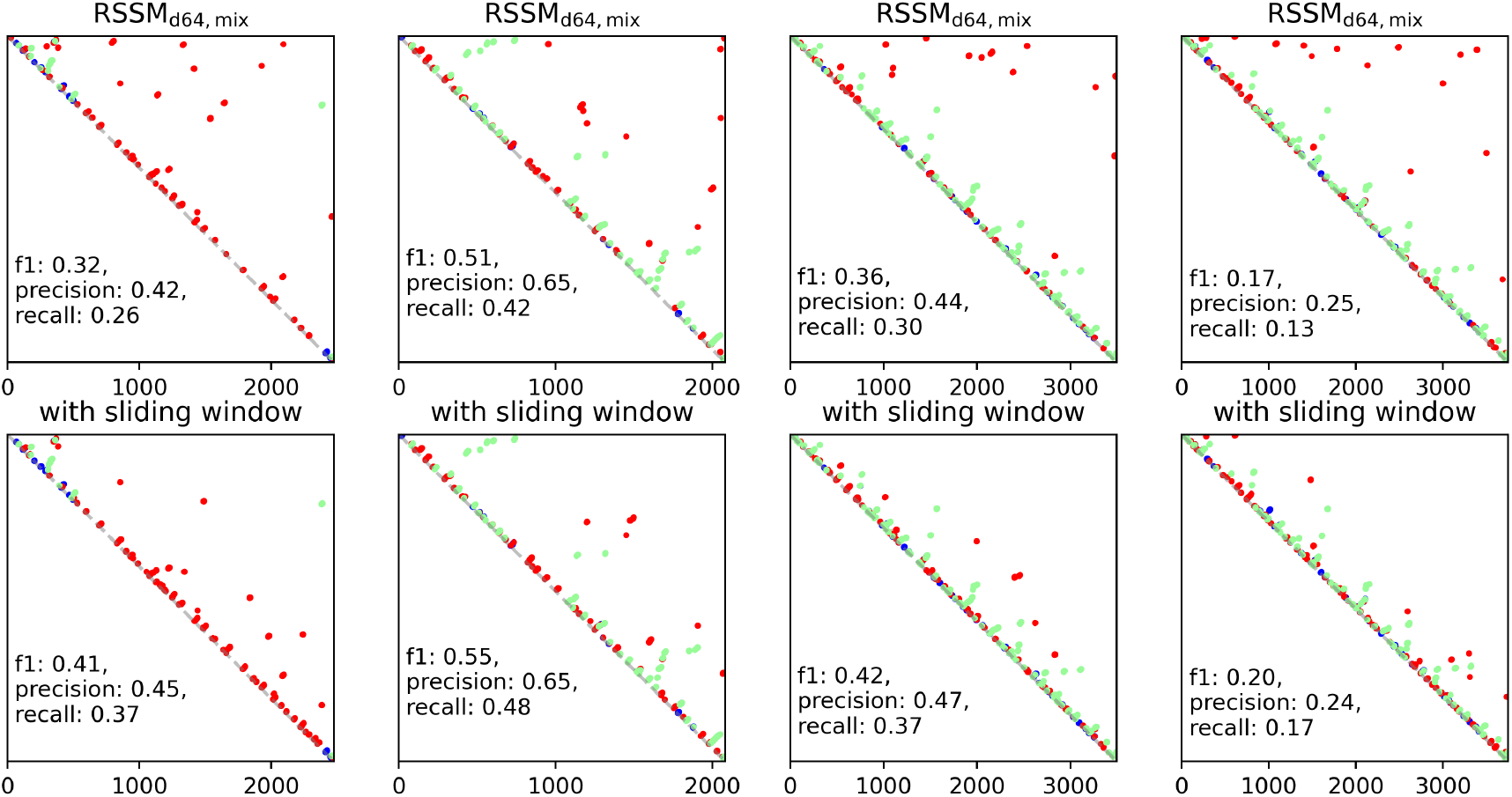
Four randomly selected long RNA examples from bprna-1m dataset. Top: RNA contact maps predicted by our base predictor, i.e. RSSM_d64,mix_ ensemble model; Bottom: RSSM_d64,mix_ plus the sliding window approach.

### D D RSSM performance on high resolution RNA secondary structures

**Table S6:** Performance of RSSM series on high resolution RNA structures. Results of baselines are taken from SPOT-RNA paper, except IPknot and ThreshKnot which are new methods introduced to these dataset.

| Model | PDB TS1 dataset (size: 67) |  |  | NMR TS2 dataset (size: 39) |  |  |
| --- | --- | --- | --- | --- | --- | --- |
|  | F1 | Precision | Recall | F1 | Precision | Recall |
| IPknot |  |  |  |  |  |  |
| LinearPartition-C | 0.612 | 0.769 | 0.524 | 0.759 | 0.918 | 0.665 |
| LinearPartition-V | 0.610 | 0.722 | 0.538 | 0.763 | 0.913 | 0.674 |
| ThreshKnot |  |  |  |  |  |  |
| LinearPartition-C | 0.604 | 0.808 | 0.499 | 0.755 | 0.926 | 0.658 |
| LinearPartition-V | 0.596 | 0.711 | 0.523 | 0.770 | 0.913 | 0.682 |
| RNAfold | 0.585 | 0.724 | 0.491 | 0.781 | 0.919 | 0.679 |
| CONTRAFold | 0.611 | 0.765 | 0.508 | 0.771 | 0.896 | 0.677 |
| Knotty | 0.603 | 0.742 | 0.508 | 0.783 | 0.913 | 0.686 |
| ContextFold | 0.621 | 0.811 | 0.503 | 0.739 | 0.910 | 0.623 |
| SPOT-RNA | 0.687 | 0.888 | 0.562 | 0.802 | 0.925 | 0.708 |
| RSSM<sub>d64</sub> | 0.656 | 0.843 | 0.550 | 0.707 | 0.853 | 0.614 |
| RSSM<sub>d64,T</sub> | 0.661 | 0.852 | 0.552 | 0.703 | 0.865 | 0.608 |
| RSSM<sub>d64,mix</sub> | 0.694 | 0.857 | 0.595 | 0.746 | 0.913 | 0.649 |
| RSSM<sub>d64,mix,LPC</sub> | 0.700 | 0.861 | 0.602 | 0.751 | 0.913 | 0.655 |

Our RSSMs have shown comparable or superior performance compared to a diverse selection of baselines in bprna-1m and Rfam derived dataset. Here, we aim to offer a glimpse of their performance on a smaller set of high resolution RNA structures obtained with techniques such as X-ray crystallography and nuclear magnetic resonance (NMR).

These high resolution RNA secondary structures are originally curated by Singh *et al*. (2019), where the proposed SPOT-RNA is initially trained on bprna training set, then finetuned on PDB TR1 set, validated on PDB VL1 set and finally tested on PDB TS1 and NMR TS2 set ^5^. Here, we skip the step of finetuning our RSSMs on the high resolution PDB TR1 and PDB VL1 set. Instead, we directly measure the performance of our RSSMs in PDB TS1 and NMR TS2, as shown in Table S6.

In both dataset of high resolution RNA structures, once again we observe benefits of pretraining RSSM with Rfam curated data, mixing diverse model hypotheses and integrating LinearPartition prediction in the ensemble, which are consistent with our observation made earlier in bprna-1m and Rfam derived dataset. In particular, having certain members in our RSSM ensemble to more thoroughly leverage the large quantity of computationally predicted RNA secondary structures also tangibly improves its overall performance on high resolution RNA secondary structures.

Compared to baselines in PDB TS1 dataset, our best performing RSSM variants such as RSSM_d64,mix_ and RSSM_d64,mix,LPC_ have outperformed SPOT-RNA without soliciting any additional supervision from high resolution RNA secondary structures. On the other hand, in NMR TS2 dataset our RSSMs are outperformed by SPOT-RNA and traditional RNA folding algorithms such as RNAfold and CONTRAfold, while such drawback does not hold statistical significance given the small amount of high resolution RNA structures in this dataset.

### E E Comparison of discretization strategies

Given a symmetric RNA contact map *M*^*p*^ ∈ ℝ^*l*×*l*^ densely populated by predicted basepairing probabilities, one last procedure is to convert this continuous representation to a discrete RNA secondary structure *Y* ∈ { 0, 1 }^*l*×*l*^. Note that *Y* by definition is required to be symmetrical, but in practice this can be relaxed by only computing the upper triangular portion of *Y*. The discretization process proceeds by maximizing the cumulative basepairing probabilities above a certain threshold *t*, i.e. 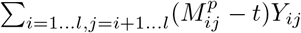, while observing a set of constraints that governs the space of admissible RNA secondary structures, which usually are:

- no overlapping of basepairs (no multiplets), i.e. Σ_*j*=*i*+1…*l*_ *Y*_*i*·_ ∈ {0, 1}, ∀*i* = 1 … *l*
- no sharp hairpin loops, i.e. ∀ *Y*_*ij*_ = 1, |*i* − *j*| ≥ 4
- only canonical pairs or wobble pairs, i.e. ∀ *Y*_*ij*_ = 1, *x*_*i*_*x*_*j*_ ∈ {*AU, UA, CG, GC, GU, UG*}

In this study, we only consider the first two types of constraints, which allows our RSSM to enrich an RNA secondary structure with any non-canonical interactions. Our default discretization strategy, which is used to report all performance in this paper unless otherwise specified, is a greedy approach that iterates 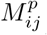 above the threshold from highest to lowest and tries to form as many basepairs as possible without violating the constraints. Our default threshold is set to 0.5. We have also considered alternative strategies based on beam search which keep multiple trajectories of sampled pairs that are no longer greedily selected.

Throughout the lines of existing work, we identified some popular discretization methods, such as the one used by SPOT-RNA and SPOT-RNA2, which we refer to as conflict sampling since it essentially reduces multiplet clusters to nonoverlapping pairs. We also include the postprocessing technique known as differentiable linear programming which is employed by E2Efold ^6^, and the ProbKnot approach from ThreshKnot. Finally, we include a naive approach referred as no sampling, which basically pairs any two bases above the threshold while disregarding the constraints. We compare all these approaches together in Figure S5 using our RSSM_d64,mix_ on the Rfam derived dataset and with varying thresholds.

As shown in figure S5, ProbKnot and conflict sampling perform better when we use a threshold below 0.1. Beam search based methods perform better on short and medium RNAs when the threshold varies between 0.2 and 0.4, while the difference between these two are not significant and the gain over our default greedy method is fairly marginal. The global maximum of F1 performance is usually attained at a threshold between 0.2 and 0.3.

Shuffled beam search and the naive no-sampling methods are in particular susceptible to low threshold values. However, as the threshold grows larger than 0.6, pairing as many basepairs while overlooking the constraints as in the no-sampling approach begins emitting marginally better performance, since fewer positive predictions remain after filtering through a larger threshold. At this point, F1 performance is mainly influenced by small variation in recall, since precision has become high but largely static. For longer RNAs without pseudoknots, the global maximum is markedly obtained by the naive approach, due to the large quantity of low basepairing probabilities predicted by RSSM as it is uncertain on these examples. In this case, variations in the recall also influence more on the harmonized F1 value.

Overall, we have shown that basepairing probabilities predicted by RSSM are amenable to a wide variety of discretization techniques, some of which are implemented in this work while others come from previous studies. In Rfam derived dataset, most discretization methods have performed comparably well on low threshold values, with the best overall performance usually obtained at a threshold between 0.2 and 0.3.

**Figure S5:**
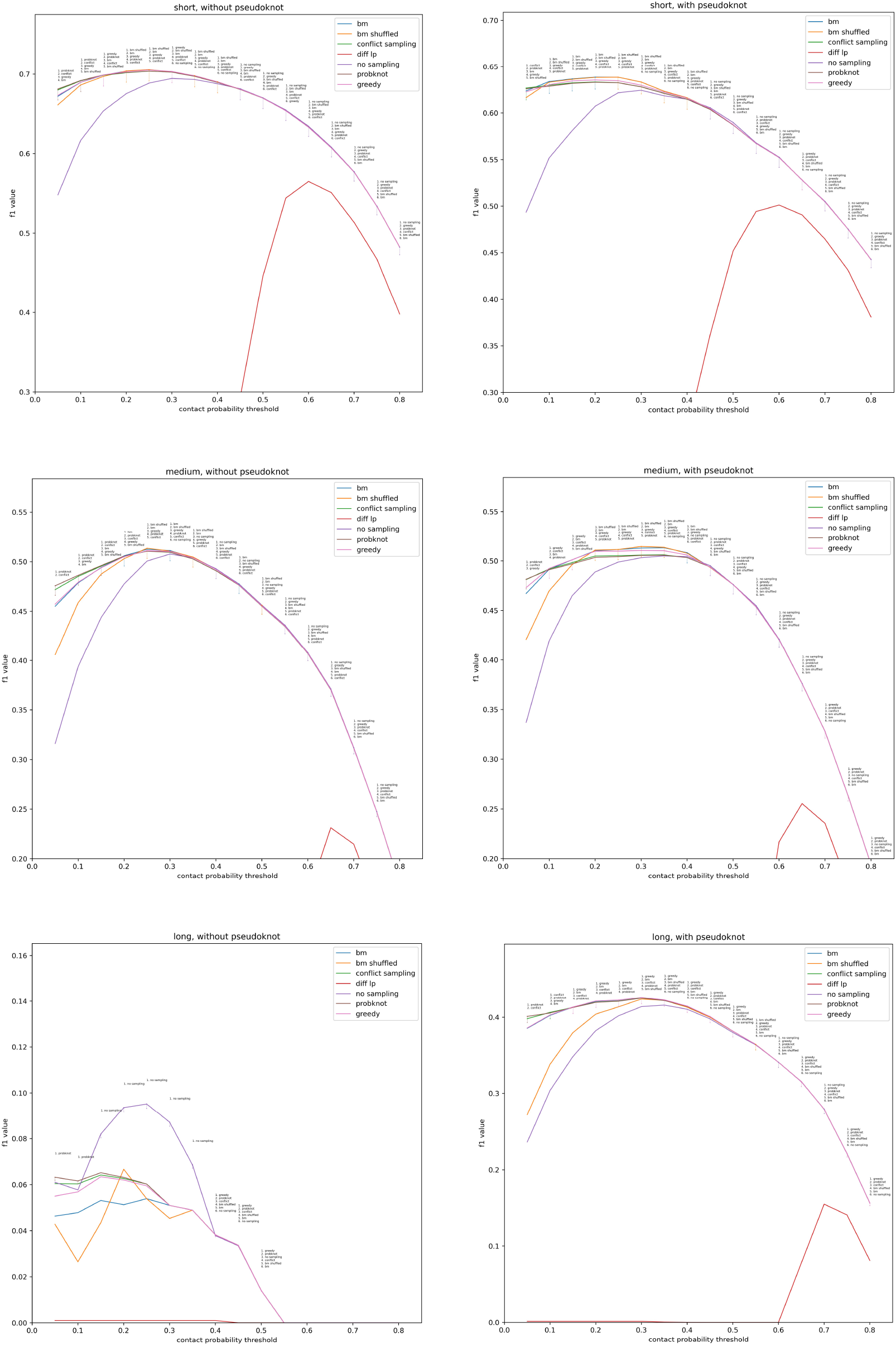
bm and bm shuffled stand for two beam search based discretization methods — the first one iterates basepairing probabilities sorted from highest to lowest, while the second one proceeds by randomly shuffled basepairing probabilities. diff lp stands for differential linear programming that is used by E2Efold. We compare all these methods across different RNA types in the Rfam derived dataset with varying thresholds. Methods that have less than 2% performance difference from the best at each threshold are ranked and marked. Zoom in on these figures is suggested.

### F F Effect of model ensembling

Figure S6 and S7 show the perplexity of basepairing probabilities averaged within the upper triangular portion of predicted RNA contact maps, either from the ensemble model or one of its five replicates, versus length of the RNA. Each point indicates an RNA sequence in the Rfam derived dataset.

In every case, there is positive correlation between the length of RNA and the averaged perplexity, which means determining basepairs becomes more difficult as the length of RNA increases. However, such positive correlation in our ensemble model is much smaller compared to any of its replicate, which indicates that ensemble averaged RNA contact maps tend to ease the difficulty of sampling basepairs.

Together with our observation made in Figure S8, the benefit of model ensembling may possibly be explained by the trimming of outlier predictions that have high probabilities in only a few replicate models, while retaining more confident basepairs that are agreed upon by most replicates inside the ensemble.

**Figure S6:**
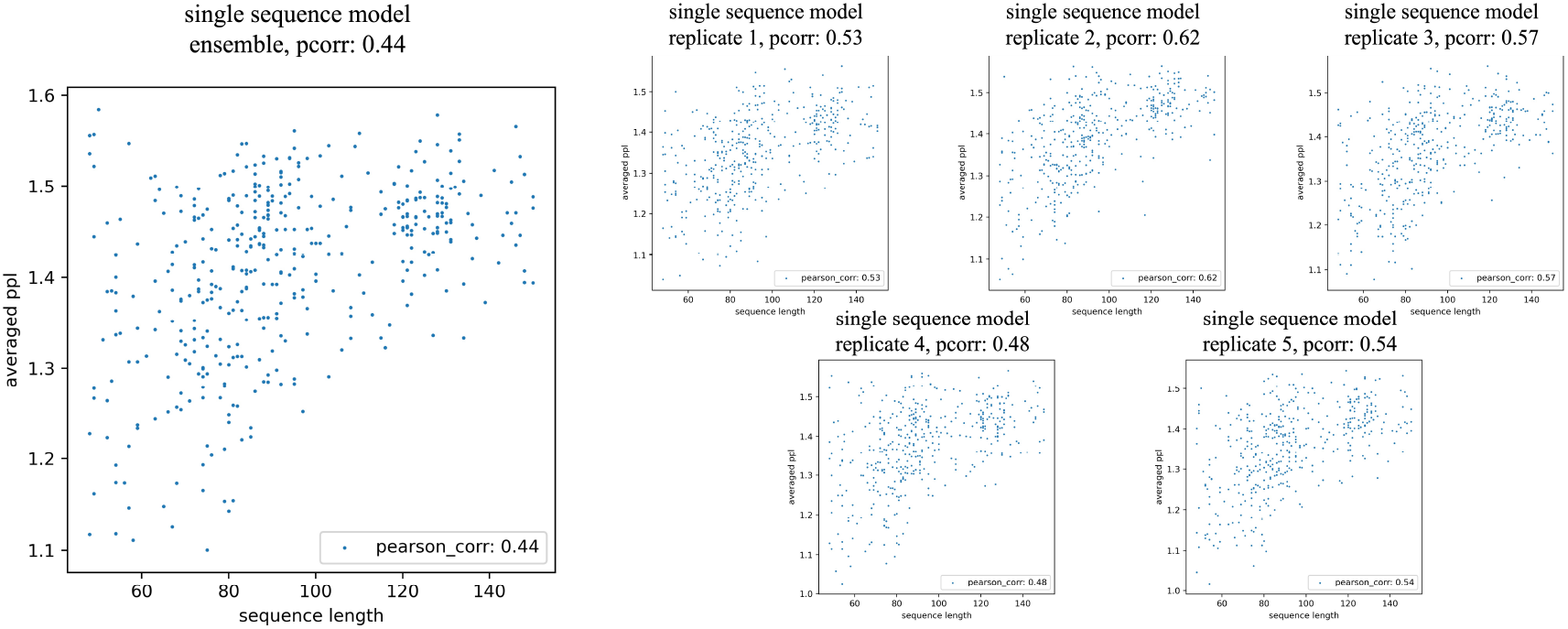
Using short RNA sequences without pseudoknot in the Rfam derived dataset. Each point indicates an RNA sequence.

**Figure S7:**
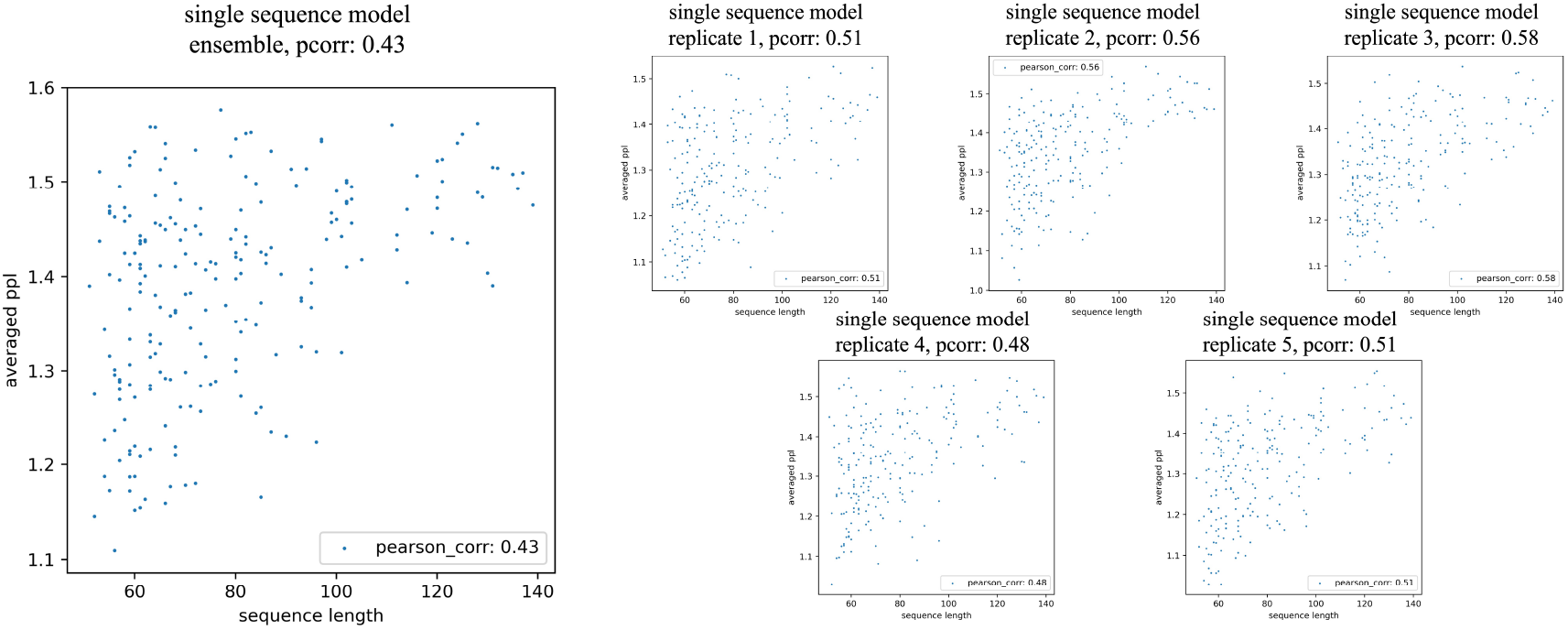
Using short RNA sequences with pseudoknots in the Rfam derived dataset. Each point indicates an RNA sequence.

**Figure S8:**
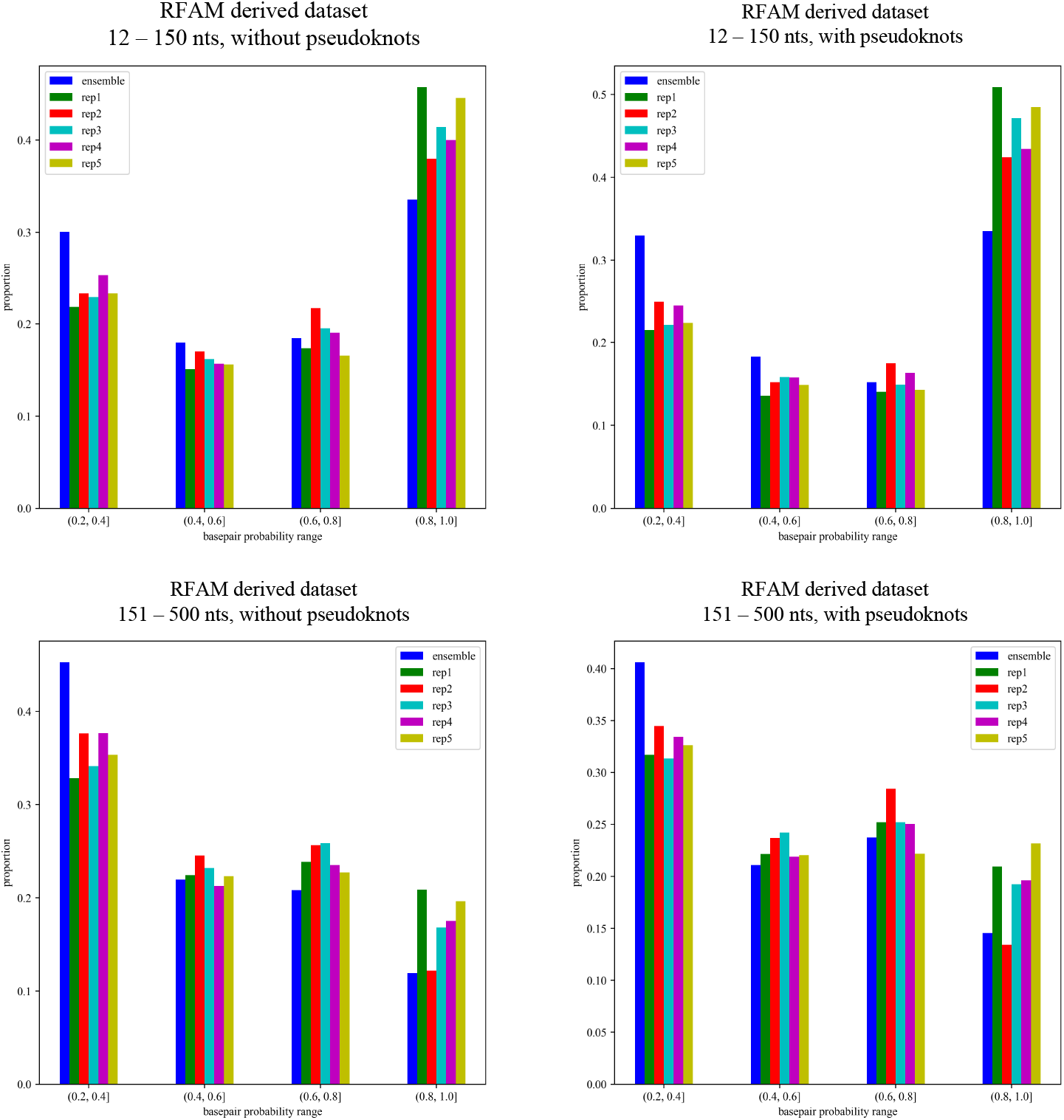
Proportion of predicted basepairing probabilities that lie within ranges such as (0.2, 0.4], (0.4, 0.6], (0.6, 0.8] and (0.8, 1.0], for RNAs in the Rfam derived dataset from short to medium length, with or without pseudoknots. Comparing ensemble averaged basepairing probabilities to those from single models, a general trend that is a reduction of large probabilities while accompanied by an increase in low probabilities is consistently observed across different RNA types.

### G G Transformer overfitting

**Figure S9:**
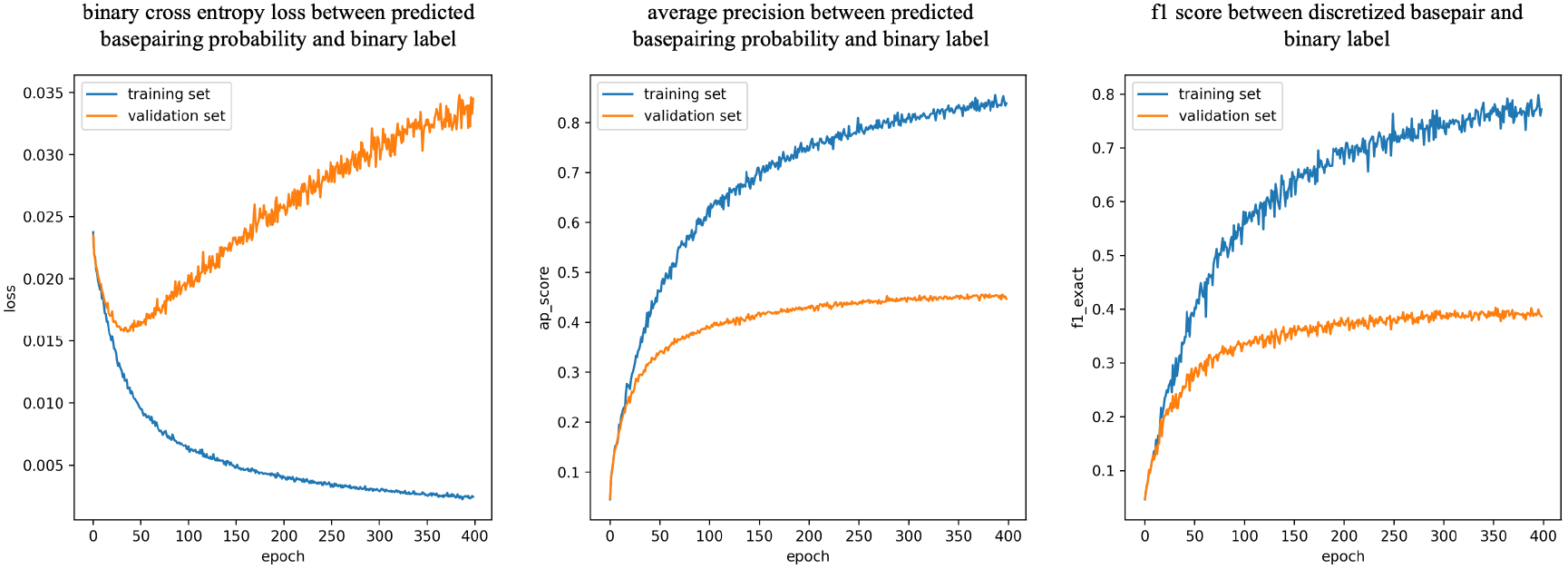
Training a simple transformer based model leads to severe overfitting in bprna dataset. This model contains five standard transformer encoder layers with base dimension of 64 and four self attention heads in each layer. Input RNA sequences are first embedded into vectors of dimension 64, with addition of learnable positional embeddings, before going through these transformer layers. Output from transformer layers are outer-concatenated to form contact maps (dimension 128), and basepairing probabilities are finally obtained with an MLP — linear(128, 128), ReLU and linear(128, 1). Dropout of 0.1 is used in transformer layers. Adam optimizer with a flat learning rate of 1e-4 is used to train the entire model. Figure clearly shows that a small transformer based model can still severely overfit the data, and the trend is only apparent when the overlap between training and validation set is properly removed as bprna dataset has ensured.

### H H Comparison of evolutionary information

**Figure S10:**
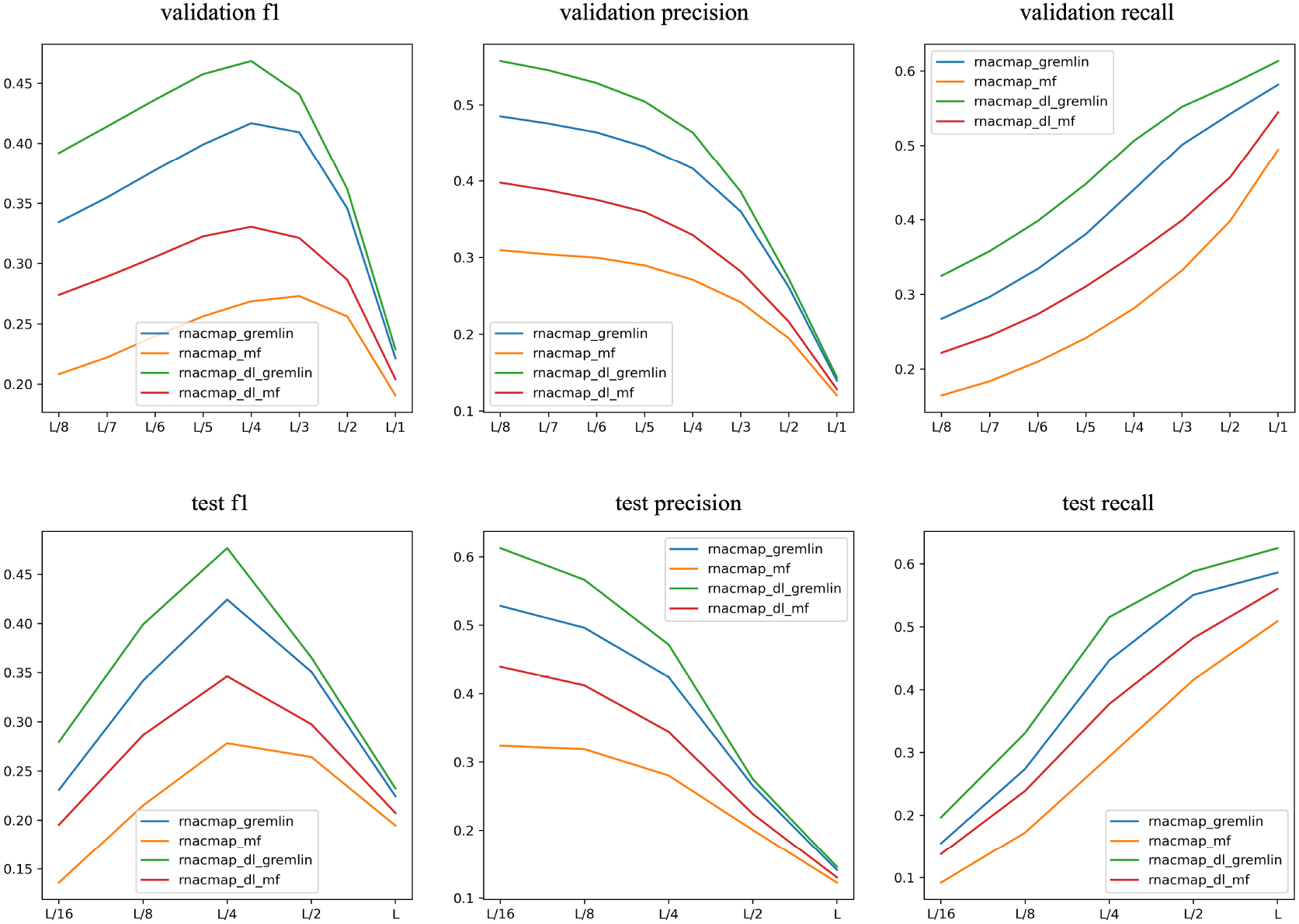
A comparison between two types of MSA with consensus secondary secondary structure either annotated by RNAfold (denoted by rnacmap) or one RSSM_d64,T_ replicate (denoted by rnacmap_dl). Performance is measured on bprna validation and test set.

We extract direct coupling signals directly from those two types of MSA using either mean-field DCA (mfDCA) or pseudo-likelihood based DCA (known as gremlin). The strength of coupling between any two MSA columns, which may indicate basepairing given the fact that paired bases in RNA structures usually undergo evolutionary co-conservation, is a scalar measured by the norm of DCA parameters followed by average product correction (APC) Morcos *et al*. (2011). Then, we rank these coupling scores from highest to lowest, and indiscriminately selects the top L/16, L/8, L/4, L/2 or L as basepairs.

These figures clearly favour evolutionary sources with underlying consensus secondary structures annotated by our own deep learning tool instead of RNAfold. Structures annotated by our tools are in general more accurate than those from RNAfold, which may help to build more accurate covariance models inside the RNAcmap pipeline and recover more pertinent sequences from the NCBI nucleotide database that tend to be truly evolutionarily related homologs.

### I I Full performance of alignment based models in bprna dataset

**Table S7:** RSSM with evolutionary features compared to a variety of baselines. Setup is similar to single sequence based RSSM, except all alignment based models are trained 500 epochs. Subscript also indicates the type of consensus secondary structure annotated inside the RNAcmap pipeline, which leads to different MSAs hence consequently affecting the performance of our alignment based RSSMs.

| Mode | Model | Validation set |  |  | Test set |  |  |
| --- | --- | --- | --- | --- | --- | --- | --- |
|  |  | F1 | Precision | Recall | F1 | Precision | Recall |
| Single models | RSSM<sub>d64,rnafold</sub> | 0.737±0.003 | 0.746±0.003 | 0.746±0.004 | 0.733±0.003 | 0.742±0.002 | 0.742±0.004 |
|  | RSSM<sub>d64,T,rnafold</sub> | 0.745±0.002 | 0.763±0.005 | 0.748±0.005 | 0.740±0.004 | 0.755±0.009 | 0.745±0.005 |
|  | RSSM<sub>d64,dl</sub> | 0.739±0.001 | 0.760±0.002 | 0.738±0.003 | 0.742±0.001 | 0.764±0.002 | 0.738±0.002 |
|  | RSSM<sub>d64,T,dl</sub> | 0.745±0.001 | 0.767±0.004 | 0.744±0.003 | 0.747±0.002 | 0.769±0.002 | 0.743±0.004 |
| Ensemble models | RSSM<sub>d64,rnafold</sub> | 0.750 | 0.769 | 0.752 | 0.746 | 0.766 | 0.747 |
|  | RSSM<sub>d64,T,rnafold</sub> | 0.758 | 0.788 | 0.754 | 0.752 | 0.779 | 0.749 |
|  | RSSM<sub>d64,dl</sub> | 0.747 | 0.776 | 0.741 | 0.749 | 0.779 | 0.741 |
|  | RSSM<sub>d64,T,dl</sub> | 0.752 | 0.781 | 0.747 | 0.754 | 0.782 | 0.747 |
| End-to-end models | MSA-RSSM<sub>d64,T,rnafold</sub> | 0.782 | 0.796 | 0.786 | 0.777 | 0.793 | 0.776 |
| Baselines | SPOT-RNA2-IL0 | 0.727 | 0.731 | 0.724 | 0.727 | 0.731 | 0.723 |
|  | SPOT-RNA2-IL1 | 0.738 | 0.779 | 0.701 | 0.734 | 0.780 | 0.693 |
|  | SPOT-RNA2-IL2 | 0.726 | 0.714 | 0.738 | 0.726 | 0.716 | 0.737 |
|  | RNAalifold (MEA) | 0.548 | 0.687 | 0.496 | 0.552 | 0.695 | 0.498 |
|  | CentroidAlifold (MEA, noncanonical) | 0.623 | 0.622 | 0.680 | 0.626 | 0.629 | 0.678 |
|  | IPknot (LinearPartition-C) | 0.632 | 0.683 | 0.626 | 0.633 | 0.687 | 0.627 |

Here, we present the full result of our alignment based RSSMs. All our single and ensemble models are trained under similar settings as single sequence based models in Table 2, except all our RSSMs are trained 500 epochs, in addition to the integration of evolutionary information either in the form of extracted covariance feature or learnt end-to-end by MSA transformer.

The benefits of Rfam pretraining and model ensembling, once again, can be confirmed in Table S7. Different types of evolutionary sources that are MSAs with inherent consensus secondary structures either annotated by our own deep learning tool (i.e. one RSSM_d64,T_ replicate) or RNAfold inside the RNAcmap pipeline also have an tangible impact on RSSM performance, and we will elaborate further on that effect in section J. We also included a few representative baselines that are SPOT-RNA2, RNAalifold, CentroidAlifold and IPknot.

SPOT-RNA2 is an alignment based deep learning model that integrates intermediate features extracted from a range of existing bioinformatics tools, such as basepairing probabilities from LinearPartition and pseudo-likelihood based DCA scores from gremlin, in addition to the position specific scoring matrix (PSSM) extracted from the original MSA. SPOT-RNA2-IL[0-2] represents three model architectures that have been directly trained on the bprna dataset, using MSAs annotated by SPOT-RNA which is an earlier single sequence based deep learning model as shown in Table 2.

We are able to show that by only using covariance features and MSAs annotated by RNAfold, our RSSM_d64,rnafold_ can obtain comparable performance to the best SPOT-RNA2 variant, and we will show later in section J how this RSSM configuration is more robust to potential perturbations in MSAs. Rfam pretraining, model ensembling and using MSAs annotated by our own deep learning model have further consolidated the superiority of our RSSMs over SPOT-RNA2.

As inputs to traditional alignment based RNA folding algorithms such as RNAalifold, CentroidAlifold and IPknot (switched to alignment based consensus structure prediction mode), we extract a representative set of 2000 sequences sampled from each RSSM_d64,T_ annotated MSA, since deeper MSAs can incur prohibitively long run time. For example, under the current setting, it takes CentroidAlifold 38 CPU hours and IPknot 22 CPU hours to go over the entire bprna validation set which contains 1300 RNA sequences. Configurations of these tools are indicated inside parentheses. Once again, we have demonstrated the superiority of representation learning enabled approaches such as RSSMs and SPOT-RNA2 over traditional approaches in alignment based RNA secondary structure prediction paradigm.

In order to better integrate evolutionary information from MSAs to our RSSMs, we leverage a dedicated neural architecture that has been introduced as MSA transformer, which we have properly pretrained using Rfam deposited MSAs as part of our pretraining strategies delineated in section “Pretraining strategies”. During the course of finetuning in bprna dataset and on MSAs obtained with RNAcmap, as usual we outer-concatenate the representation of the target RNA sequence within the learnt MSA and extract all row-wise attention weights, which are then concatenated together and provided to RSSM as input that carries evolutionary signals in place of covariance features.

This strategy of combining MSA transformer and RSSM enables us to obtain a fully end-to-end deep learning model referred as MSA-RSSM, which is similar to how MSA transformer was originally used to perform supervised protein contact map prediction in Rao *et al*., 2021.

During the course of finetuning MSA-RSSM, we set the maximal length of randomly subsampled MSA depth at *m* = 2^13^*/l*, for the sake of reducing computational overhead and improve training efficiency. At the inference stage, however, the type of MSA subsampling strategy can generate a large impact on the final performance of MSA-RSSM. Rao *et al*., 2021 originally showcased a diversity maximizing subsampling strategy and cautioned that subsampling larger MSA than what has been seen during training may degrade model performance, which we are able to confirm in Figure S11 A.

However, considering the fact that randomly subsampled MSAs may well induce comparable performance as mentioned in Rao *et al*., 2021 and again confirmed in Figure S11 B, and in order to fully tap into the rich evolutionary information covered by a potentially large MSA, we once again borrow the idea of ensemble averaging multiple RNA contact maps, but this time instead of soliciting contact prediction from different models, we stick to a single MSA-RSSM and each contact map is predicted from a randomly subsampled MSA.

In practice, for each MSA with original depth larger than the threshold *m* = 2^13^*/l*, we randomly subsample it 10 times and ensemble average all the predicted RNA contact maps. Considering the stochastic nature of random MSA subsampling, we repeat this process 10 times over the entire dataset and register the mean performance in Table S7, while the standard deviation is reflected as shaded area in Figure S11 A. We use this MSA subsampling strategy to report MSA-RSSM performance in Table S7, and we are able to confirm its superiority over the diversity maximizing principle in Figure S11 A.

### J J Robustness of alignment based models

We study the robustness of our alignment based models from the following two perspectives:

1. The effect of evaluating properly trained RSSMs on a different type of MSA with underlying consensus secondary structure annotated by a different tool from the one used in training.
2. The influence of MSA depth on the performance of our RSSMs.

#### J.1 J.1 Swapping evolutionary sources

The logic behind the first angle is to understand how a properly finetuned alignment based RSSM would behave when it is given a different type of MSAs from those the model has actually seen during training, since there are no constraints on the type of evolutionary information that our models might receive in practice. Here, we only focus on the effect of different forms of consensus secondary structure annotation, although other influencing factors are also worth considering, such as MSAs with no consensus secondary structures at all (e.g. blastn seed alignment) similar to those used in protein structure prediction, which we will have to leave for future investigation.

We present the result in Table S8, which are experiments of swapping input evolutionary sources at the inference stage. Several important observations can be made. First, we notice that RSSMs trained with MSAs annotated by RNAfold have all performed better on RSSM_d64,T_ annotated MSAs, and the performance gain is consistent across our single models, ensemble models as well as end-to-end MSA-RSSM model. However, RSSMs trained with deep learning annotated MSAs are conspicuously susceptible to the other type of MSAs, for which we hope to offer a possible explanation below.

**Table S8:** The performance of alignment based models using different types of evolutionary sources (MSAs). For example, RSSM_d64,T,rnafold←dl_ means an RSSM is trained using MSAs with underlying consensus secondary structures annotated by RNAfold, while the performance is reported with MSAs annotated by our own deep learning tool.

| Mode | Model | Validation set |  |  | Test set |  |  |
| --- | --- | --- | --- | --- | --- | --- | --- |
|  |  | F1 | Precision | Recall | F1 | Precision | Recall |
| Single models | $\text{RSSM}_{d64,\text{rnafold}}$ | $0.737 \pm 0.003$ | $0.746 \pm 0.003$ | $0.746 \pm 0.004$ | $0.733 \pm 0.003$ | $0.742 \pm 0.002$ | $0.742 \pm 0.004$ |
| | $\text{RSSM}_{d64,\text{rnafold} \leftarrow \text{dl}}$ | $0.752 \pm 0.002$ | $0.771 \pm 0.002$ | $0.752 \pm 0.003$ | $0.753 \pm 0.001$ | $0.774 \pm 0.002$ | $0.751 \pm 0.002$ |
| | $\text{RSSM}_{d64,T,\text{rnafold}}$ | $0.745 \pm 0.002$ | $0.763 \pm 0.005$ | $0.748 \pm 0.005$ | $0.740 \pm 0.004$ | $0.755 \pm 0.009$ | $0.745 \pm 0.005$ |
| | $\text{RSSM}_{d64,T,\text{rnafold} \leftarrow \text{dl}}$ | $0.755 \pm 0.005$ | $0.778 \pm 0.004$ | $0.753 \pm 0.008$ | $0.754 \pm 0.003$ | $0.778 \pm 0.005$ | $0.751 \pm 0.006$ |
| | $\text{RSSM}_{d64,\text{dl}}$ | $0.739 \pm 0.001$ | $0.760 \pm 0.002$ | $0.738 \pm 0.003$ | $0.742 \pm 0.001$ | $0.764 \pm 0.002$ | $0.738 \pm 0.002$ |
| | $\text{RSSM}_{d64,\text{dl} \leftarrow \text{rnafold}}$ | $0.645 \pm 0.002$ | $0.622 \pm 0.005$ | $0.699 \pm 0.003$ | $0.643 \pm 0.003$ | $0.623 \pm 0.005$ | $0.691 \pm 0.004$ |
| | $\text{RSSM}_{d64,T,\text{dl}}$ | $0.745 \pm 0.001$ | $0.767 \pm 0.004$ | $0.744 \pm 0.003$ | $0.747 \pm 0.002$ | $0.769 \pm 0.002$ | $0.743 \pm 0.004$ |
| | $\text{RSSM}_{d64,T,\text{dl} \leftarrow \text{rnafold}}$ | $0.645 \pm 0.002$ | $0.617 \pm 0.004$ | $0.707 \pm 0.002$ | $0.644 \pm 0.002$ | $0.620 \pm 0.004$ | $0.701 \pm 0.003$ |
| Ensemble models | $\text{RSSM}_{d64,\text{rnafold}}$ | 0.750 | 0.769 | 0.752 | 0.746 | 0.766 | 0.747 |
| | $\text{RSSM}_{d64,\text{rnafold} \leftarrow \text{dl}}$ | 0.763 | 0.793 | 0.756 | 0.765 | 0.799 | 0.754 |
| | $\text{RSSM}_{d64,T,\text{rnafold}}$ | 0.758 | 0.788 | 0.754 | 0.752 | 0.779 | 0.749 |
| | $\text{RSSM}_{d64,T,\text{rnafold} \leftarrow \text{dl}}$ | 0.766 | 0.800 | 0.759 | 0.766 | 0.802 | 0.755 |
| | $\text{RSSM}_{d64,\text{dl}}$ | 0.747 | 0.776 | 0.741 | 0.749 | 0.779 | 0.741 |
| | $\text{RSSM}_{d64,\text{dl} \leftarrow \text{rnafold}}$ | 0.652 | 0.630 | 0.704 | 0.648 | 0.628 | 0.695 |
| | $\text{RSSM}_{d64,T,\text{dl}}$ | 0.752 | 0.781 | 0.747 | 0.754 | 0.782 | 0.747 |
| | $\text{RSSM}_{d64,T,\text{dl} \leftarrow \text{rnafold}}$ | 0.650 | 0.625 | 0.712 | 0.649 | 0.626 | 0.703 |
| End-to-end models | $\text{MSA-RSSM}_{d64,T,\text{rnafold}}$ | 0.782 | 0.796 | 0.786 | 0.777 | 0.793 | 0.776 |
| | $\text{MSA-RSSM}_{d64,T,\text{rnafold} \leftarrow \text{dl}}$ | 0.786 | 0.604 | 0.787 | 0.790 | 0.809 | 0.786 |
| SPOT-RNA2 Ensemble | $\text{SPOT-RNA2}_{\leftarrow \text{rnafold}}$ | 0.546 | 0.479 | 0.676 | 0.550 | 0.485 | 0.676 |
| | $\text{SPOT-RNA2}_{\leftarrow \text{dl}}$ | 0.617 | 0.560 | 0.726 | 0.624 | 0.566 | 0.733 |

Considering our previous observation in Figure S10 which establishes that RSSM_d64,T_ annotated MSAs contain more useful co-conservation signals that reveal potential basepairs, RSSMs trained with deep learning annotated MSAs may become overly optimistic and trustful on the covariance feature and such bias does not fare well when it meets a weaker form of evolutionary sources. Indeed, we have observed that RSSMs trained with RSSM_d64,T_ annotated MSAs converged much faster than those using RNAfold annotated MSAs, which points to potential overfitting. On the other hand, when RSSMs are trained using a weaker evolutionary source that is annotated by RNAfold, our models may learn to become aware of its limitation and alert to potential spurious co-conservation signals that lie within.

Similar observation can be made for SPOT-RNA2 by evaluating this baseline on our own RSSM_d64,T_ annotated MSAs which leads to SPOT-RNA_←dl_, as well as on MSAs annotated by RNAfold that produces SPOT-RNA_←rnafold_. SPOT-RNA2 was originally trained using SPOT-RNA annotated MSAs, which is another deep learning model that has outperformed RNAfold as we have shown earlier in Table 2. Therefore, SPOT-RNA2 has similarly demonstrated a lack of robustness due to the type of evolutionary source it had used for training originally.

In summary, experiments in Table S8 suggest that for the purpose of improving the robustness of alignment based RSSMs or any other deep learning tools in general, a weaker form of MSAs should be employed for training, which may help preserve the robustness of trained models on other type of MSAs. In particular, better performance may be achieved by leveraging MSAs annotated with a more powerful form of consensus secondary structure.

Additionally, we would like to comment on the robustness of SPOT-RNA2 which may reflect our point made earlier in the Introduction about frailty in software dependency. SPOT-RNA2 requires certain types of intermediate features as input, such as basepairing probability from LinearPartition and direct coupling scores from gremlin. We have prepared input to SPOT-RNA2 exactly in the same format, except we extract direct coupling scores from a recent deep learning based gremlin implementation^7^ which runs substantially faster than the old gremlin release. SPOT-RNA2, on the other hand, is trained with the old gremlin release and it seems the performance of SPOT-RNA2 can be seriously perturbed by small nuances in the gremlin implementation. Since gremlin is a pseudo-likelihood based DCA method and its deep learning based implementation is gaining more traction, SPOT-RNA2’s dependency on its older release will become more problematic as the old gremlin release is being replaced by the new one.

**Figure S11:**
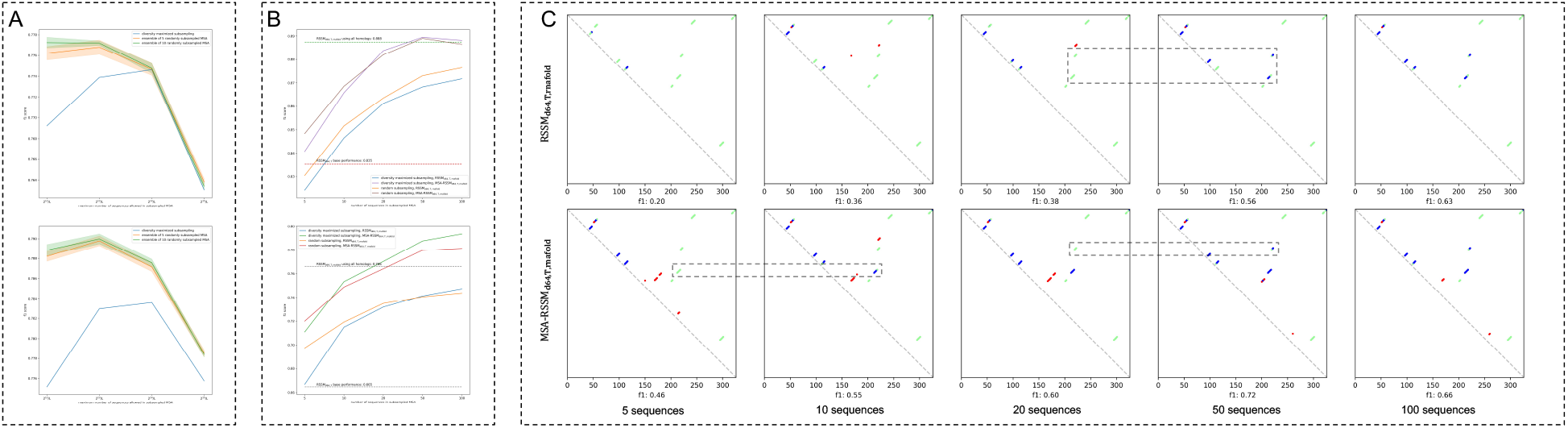
(A) Evaluating MSA-RSSM_d64,T,rnafold_ in bprna test set and comparing different MSA subsampling strategies at various maximal MSA depth thresholds; top, using RNAfold annotated MSAs; bottom, using RSSM_d64,T_ annotated MSAs. (B) Effect of MSA depth on covariance feature based RSSM_d64,T,rnafold_ and end-to-end MSA-RSSM_d64,T,rnafold_, using RSSM_d64,T_ annotated MSAs; top, high resources setting where the original MSA *N*_eff_ is larger than 1000; bottom, low resources setting where the original MSA *N*_eff_ lies between 100 and 200. (C) Visualization of predicted RNA contact maps by RSSMs using MSAs at various depth. The same set of sequences (diversity maximized) are used at each column. Blue, green and red points denote true positive, false negative and false positive basepairs respectively.

#### J.2 J.2 Effect of MSA depth on RSSM_d64,T,rnafold_ and MSA-RSSM_d64,T,rnafold_

The number of homologs found by RNAcmap can vary as it depends on the size of nucleotide database which is usually several hundreds of gigabytes (e.g. 560 GB for the NCBI database) and has been constantly growing. Therefore, in reality one may end up with substantially fewer homologs when hosting larger nucleotide database proves infeasible given a small computational budget. Besides, one may also wish to stop scanning the nucleotide database after a sufficient amount of homologs has been found, since running RNAcmap over the entire database is a certain time costly and resources intensive process.

Here, we seek to better understand how MSA depth affects our alignment based ensemble models that are RSSM_d64,T,rnafold_ and MSA-RSSM_d64,T,rnafold_, using RSSM_d64,T_ annotated MSAs. To properly simulate the scenario where we are limited to a small amount of homologous sequences, we identify two subsets of RNA sequences from bprna test set to form the test bed for: (1) a high resources setting where the original MSA *N*_eff_ is larger than 1000, and (2) a low resource setting where the original MSA *N*_eff_ is between 100 and 200. A total of 259 RNA sequences is included in the high resources setting and 157 RNA sequences is featured in the low resources setting. We gradually increase the number of homologs for each RNA, either diversity maximized or randomly sampled ^8^ from the original MSA, and show the performance of our models in Figure S11 B.

Both our alignment based models have outperformed the single sequence based ensemble baseline RSSM_d64,T_ using only 5 sequences in the low resources setting. Under the high resources setting, RSSM_d64,T,rnafold_ slightly underperforms when MSA only contains 5 sequences but the deficit is not significant. Both RSSM_d64,T,rnafold_ and MSA-RSSM_d64,T,rnafold_ benefit substantially from leveraging more RNA homologs in the MSA, as we gradually increase the MSA depth from 5 to 50. In particular, MSA-RSSM_d64,T,rnafold_ outperforms RSSM_d64,T,rnafold_ with the same amount of evolutionary information and exceeds the bar set by RSSM_d64,T,rnafold_ using all homologs in the original MSAs, when a sufficient amount of 50 homologous sequences has been reached.

Figure S11 (B) demonstrates that our end-to-end MSA-RSSM is able to better utilize evolutionary information than the covariance feature based RSSM. Both our alignment based models are relatively robust to low MSA depth and as a matter of fact, outperform single sequence based baseline as long as a minimal amount of 5 sequences are reached in the low resource setting, or 10 sequences in the high resource setting. Figure S11 (C) shows a particular RNA example where increasing MSA depth can significantly improve the performance as it helps to recall more long range basepairs (off-diagonal blue points).

### K K Generalization of alignment based RSSMs

**Table S9:** Performance of alignment based models on medium RNA sequences between 151 to 500 nts in Rfam derived dataset. For each RNA, MSA-RSSM subsamples its MSA 10 times and ensemble average each predicted contact maps. This process is repeated 10 times across the entire dataset to calculate the mean and standard deviation for MSA-RSSM.

|  |  | Rfam derived dataset (151-500 nts) |  |  |  |  |  |
| --- | --- | --- | --- | --- | --- | --- | --- |
|  |  | without pseudoknot (size: 169) |  |  | with pseudoknot (size: 70) |  |  |
|  | Model | F1 | Precision | Recall | F1 | Precision | Recall |
| Single<br>sequence<br>baselines | IPknot |  |  |  |  |  |  |
|  | LinearPartition-C | 0.506 | 0.453 | 0.610 | 0.503 | 0.475 | 0.549 |
|  | LinearPartition-V | 0.493 | 0.404 | 0.667 | 0.456 | 0.393 | 0.555 |
|  | ThreshKnot |  |  |  |  |  |  |
|  | LinearPartition-C | **0.512** | 0.494 | 0.572 | 0.490 | 0.503 | 0.500 |
|  | LinearPartition-V | 0.498 | 0.408 | 0.670 | 0.453 | 0.392 | 0.546 |
|  | RNAfold | 0.486 | 0.400 | 0.651 | 0.433 | 0.377 | 0.520 |
|  | CONTRAfold | 0.510 | 0.428 | 0.664 | 0.473 | 0.420 | 0.556 |
|  | RSSM<sub>d64,mix</sub> | 0.505 | 0.437 | 0.627 | **0.510** | 0.480 | 0.555 |
|  | RSSM<sub>d64,mix,LPC</sub> | **0.521** | 0.452 | 0.644 | **0.514** | 0.483 | 0.560 |
| RNAfold<br>annotated<br>MSAs | RNAalifold (MEA) | 0.413 | 0.585 | 0.373 | 0.435 | 0.538 | 0.392 |
|  | CentroidAlifold (MEA) | 0.501 | 0.503 | 0.534 | 0.447 | 0.469 | 0.454 |
|  | IPknot (LinearPartition-C) | 0.484 | 0.511 | 0.491 | 0.449 | 0.514 | 0.423 |
|  | gremlinDCA (top L/4) | 0.271 | 0.238 | 0.329 | 0.227 | 0.216 | 0.245 |
|  | mfDCA (top L/4) | 0.080 | 0.072 | 0.095 | 0.061 | 0.058 | 0.066 |
|  | RSSM<sub>d64,rnafold</sub> | 0.550 | 0.490 | 0.658 | 0.540 | 0.515 | 0.581 |
|  | RSSM<sub>d64,T,rnafold</sub> | **0.561** | 0.499 | 0.672 | **0.543** | 0.513 | 0.587 |
|  | MSA-RSSM<sub>d64,T,rnafold</sub> | **0.563±0.001** | 0.499±0.001 | 0.678±0.001 | **0.553±0.001** | 0.521±0.001 | 0.603±0.001 |
| RSSM<sub>d64,mix</sub><br>annotated<br>MSAs | RNAalifold (MEA) | 0.410 | 0.619 | 0.354 | 0.440 | 0.618 | 0.372 |
|  | CentroidAlifold (MEA) | 0.517 | 0.548 | 0.528 | 0.484 | 0.554 | 0.460 |
|  | IPknot (LinearPartition-C) | 0.493 | 0.543 | 0.482 | 0.476 | 0.566 | 0.439 |
|  | gremlinDCA (top L/4) | 0.294 | 0.256 | 0.361 | 0.267 | 0.253 | 0.287 |
|  | mfDCA (top L/4) | 0.090 | 0.080 | 0.108 | 0.094 | 0.091 | 0.099 |
|  | RSSM<sub>d64,rnafold</sub> | 0.562 | 0.509 | 0.658 | **0.561** | 0.537 | 0.595 |
|  | RSSM<sub>d64,T,rnafold</sub> | **0.578** | 0.516 | 0.690 | 0.550 | 0.525 | 0.590 |
|  | MSA-RSSM<sub>d64,T,rnafold</sub> | **0.569±0.002** | 0.502±0.003 | 0.688±0.002 | **0.572±0.001** | 0.544±0.001 | 0.617±0.001 |

**Table S10:**
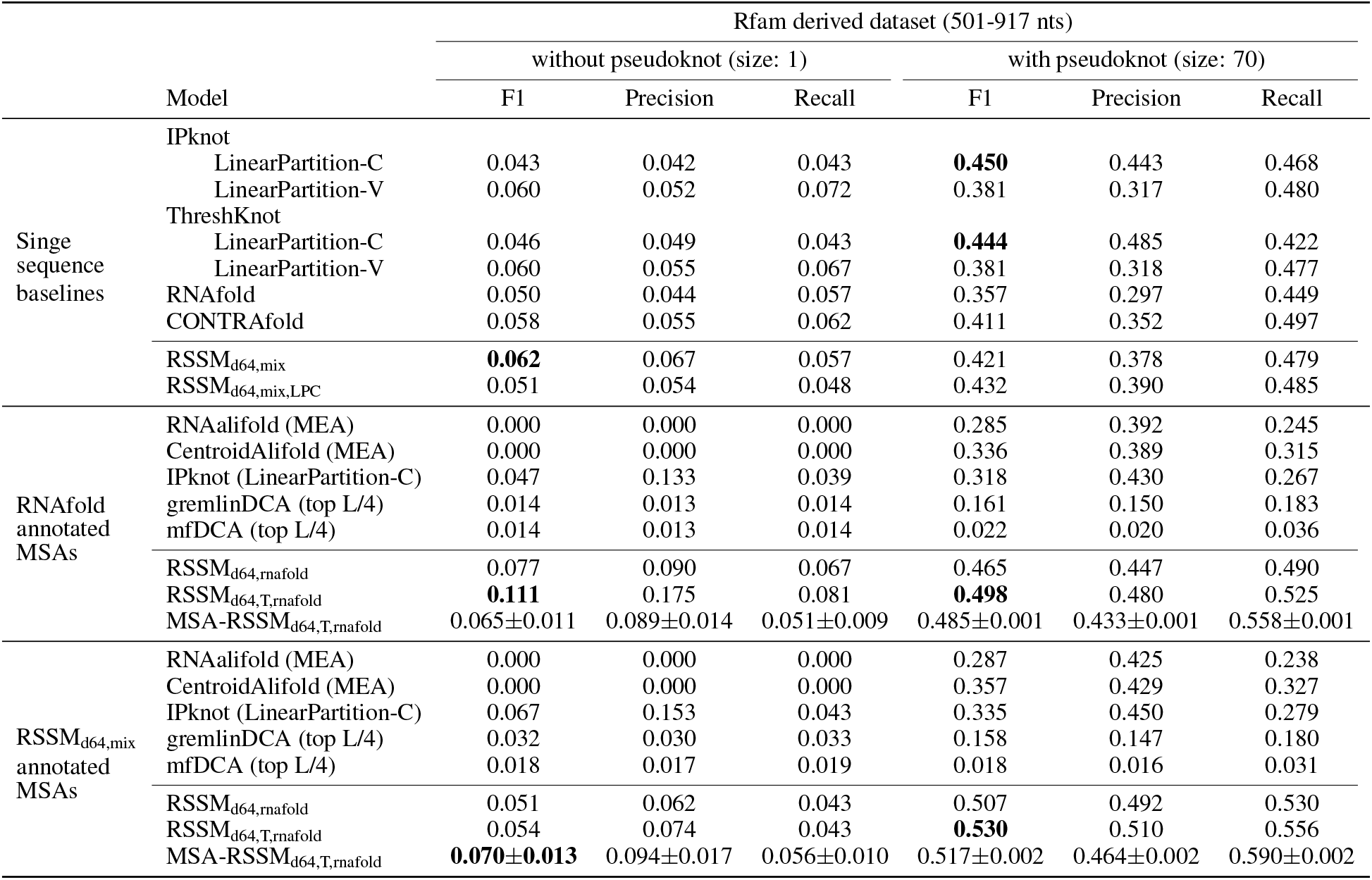
Performance of alignment based models on long RNA sequences between 501 to a maximum of 917 nts in Rfam derived dataset. For each RNA, MSA-RSSM subsamples its MSA 10 times and ensemble average each predicted contact maps. This process is repeated 10 times across the entire dataset to calculate the mean and standard deviation for MSA-RSSM.

### L L Effect of spurious homologs on prediction

**Figure S12:**
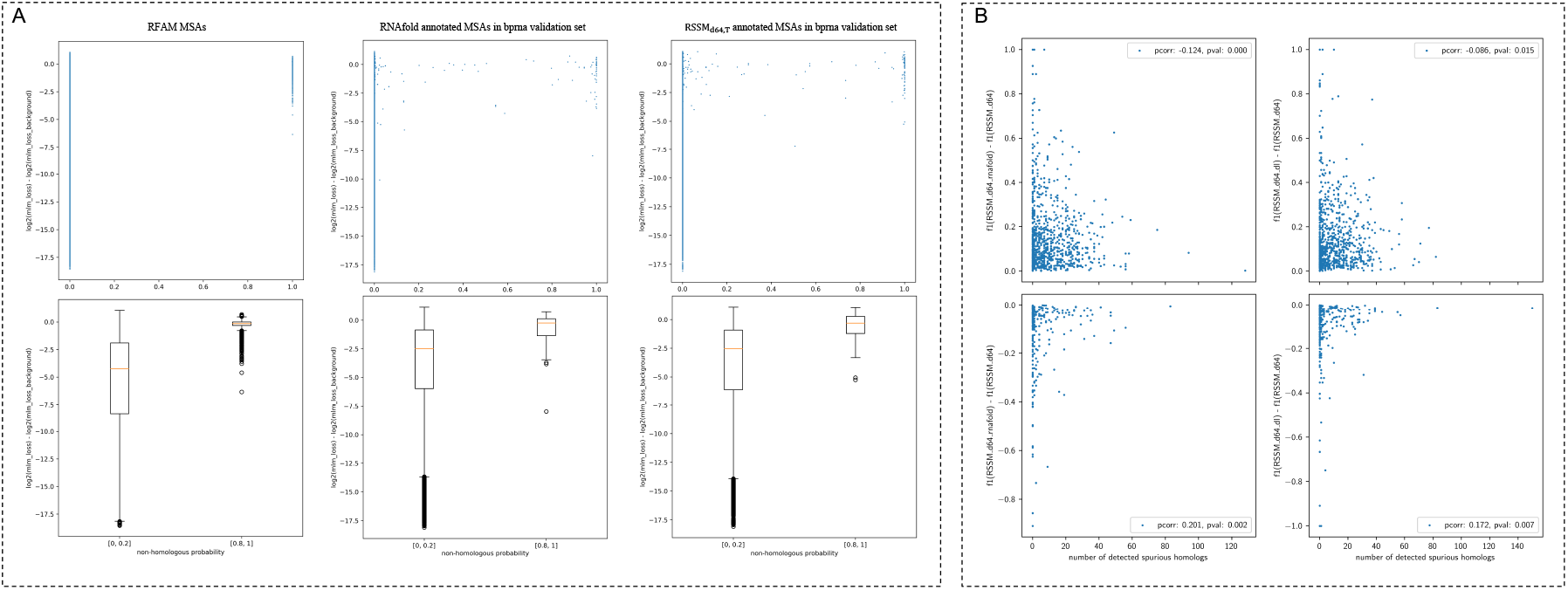
(A) Distribution of MLM loss fold change on homologs and detected non-homologs. (B) The effect of spurious homologs discovered in MSA on model performance.

Adding evolutionary information to RSSM in the form of extracted covariance features measurably improves its overall performance. However the scale of such improvement varies depending on factors such as the quality of the alignment. In particular, MSAs often include non-homologous sequences that hampered covariance-based analyses. Detecting and removing such sequences would likely improve RSSM’s performance.

Intuitively, a spurious homolog is inherently incompatible with the evolutionary information presented by a MSA, therefore reconstructing masked tokens should be difficult for those sequences. First, we first try to establish the relationship between spurious homologs that are detected by our pretrained MSA transformer and the loss of predicting masked tokens therein. However, considering the fact that reconstructing masked tokens does not always necessitate the knowledge of other homologs, where a single sequence would suffice as shown in single sequence based language models such as ESM and ProtTrans Rives *et al*. (2021); Elnaggar *et al*. (2021), we turn to measure the fold decrease of masked token prediction loss to the background loss of using only the sequence itself. Given a masked MSA input *x* ∈ ℝ^*m,l*^, the fold change of a specific masked position indexed by *i, j* is defined as:

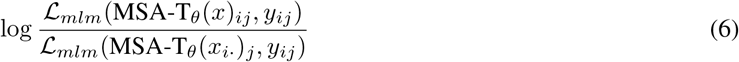

MSA-T_*θ*_ is pretrained using Rfam evolutionary data which are subsampled high quality MSA except manually contaminated noise. We discard masked tokens that can be easily inferred without other homologs by setting a threshold on the background loss, i.e. *ℒ*_*mlm*_(MSA-T_*θ*_(*x*_*i*_)_*j*_, *y*_*ij*_) *>* 5.0, and record how much fold decrease can be achieved via utilizing homologous information. We discard MSAs with less than 1000 homologs.

As shown in Figure S12 A, there is a striking difference between the fold change values obtained on masked tokens from actual homologs versus those from spurious homologs. This observation is relatively consistent across various sources of MSA. In particular, masked tokens on detected spurious homologs tend to only exhibit smaller fold decrease, which means incorporating evolutionary information from other sequences in the MSA cannot improve their prediction, hence suggesting its incompatibility with the MSA.

In Figure S12 B, we take a closer look at how potential non-homologs induce performance change. Each point is an RNA in bprna validation set and a general trend is reflected that as the number of spurious homologs increases the impact of extracted covariance features diminishes.

Overall, we are able to show a negative correlation between the amount of potential noise in a MSA and its influence on performance change. This indicates that noise within MSAs unavoidably undermine the utility of the evolutionary information it represents thereof.

## Footnotes

2 PDB dataset contains x-ray crystallography Smyth and Martin (2000) determined RNA structures from the Protein Data Bank Rose *et al*. (2017); NMR dataset contains other experimental RNA structures obtained via Nuclear Magnetic Resonance technique Le Bihan (1991).

3 We used the github release of IPknot: https://github.com/satoken/ipknot/releases, on a system with an Intel Xeon 5220R CPU

4 [See CONTRAfold manual “–noncomplementary”](http://contra.stanford.edu/contrafold/manual_v2_02.pdf)

5 PDB: Protein Data Bank; NMR: Nuclear Magnetic Resonance

6 While the method contains a larger amount of hyperparameters such as step sizes lr_min_, lr_max_, number of iterations, l_1_ coefficient *ρ* and etc., we simply adopted their original implementation on github ([link](https://github.com/ml4bio/e2efold/blob/master/e2efold/postprocess.py)).

7 [Deep learning based gremlin implementation](https://github.com/sokrypton/GREMLIN_CPP).

8 We randomly sample each MSA 10 times and produce an averaged performance. Note, in this part we do not ensemble contact maps predicted from each randomly sampled MSA.

